# Resources allocation explains the differential roles of RBS and promoter strengths in cell mass distribution and optimal protein expression productivity

**DOI:** 10.1101/2020.11.19.390583

**Authors:** Fernando Nóbel, Jesús Picó

## Abstract

Design of synthetic genetic circuits without considering the impact of host–circuit interactions results in an inefficient design process and lengthy trial-and-error iterations to appropriately tune the expression levels. Microorganisms have evolved to reach an optimal use of cellular resources. This balance is perturbed by circuit-host interactions resulting from the interaction among the cell environment from which the cell takes substrates, its metabolism, and the needs of exogenous synthetic genetic circuit introduced in the cell host. The resulting competition for common shared cell resources introduces spurious dynamics leading to problems of malfunctioning of the synthetic circuit due to lack of enough cellular resources. Therefore, there is an increasing interest in development of methods for model-based design of synthetic gene circuits considering host-circuit interactions. Here we present a small-size model of gene expression dynamics in bacterial cells accounting for host-circuit interactions. For each gene, the model defines the cellular resources recruitment strength as the key functional coefficient that allows to explain the distribution of resources among the host and the genes of interest and the relationship between the usage of resources and cell growth. This functional coefficient explicitly takes into account the availability of resources and lab-accessible gene characteristics, such as promoter and ribosome binding site (RBS) strengths and capture their interplay with the availability of free cell resources. In spite of its simplicity, the model is able to explain the differential role of promoter and RBS strengths in the distribution of protein mass and the optimal protein expression productivity with remarkable fit to the experimental data from the literature for *E. coli*. This makes the model amenable for model-based circuit design purposes. Moreover, the model also allows to understand why endogenous ribosomal and non-ribosomal genes have evolved different strategies in the expression space.

## 1. Introduction

The rational design of synthetic genetic circuits of increasing complexity, their experimental characterization and robust tuning, and their industrial scaling are some important milestones towards the integration of complex synthetic genetic circuits into microorganisms for a variety of practical applications. Within this context, one of the fundamental problems in designing synthetic genetic circuits that explain the current disparity between the ability to design biological systems and to synthesize them is the lack of systematic design methods considering the interaction with the host cell [7].

Circuit-host interactions result from the interrelations among the cell environment from which the cell takes substrates, its metabolism, and the needs of the synthetic genetic circuit introduced in the cell host. The over-expression of exogenous proteins by a genetically modified microorganism as well as the production of metabolites by the addition and/or modification of their metabolic pathways introduces a metabolic load that removes the microorganism from its natural state, which has evolved to reach an optimal use of resources [40]. The resulting competition for common shared cell resources affects cell growth and introduces spurious dynamics [31] leading to problems of malfunctioning of the synthetic circuit due to lack of enough cellular resources. It also triggers its elimination by evolutionary mechanisms trying to restore the natural optimal state [10], or both.

Design of synthetic genetic circuits without considering the impact of host–circuit interactions results in an inefficient design process and lengthy trial-and-error iterations to appropriately tune a circuit’s expression levels [43]. Therefore, in the last years the has been an increasing interest in development of methods for model-based design of synthetic gene circuits considering host-circuit interactions [33]. Indeed, burden-aware models are also useful to infer why cells have evolved particular strategies for cell expression.

The simplest burden-aware models consider the shared resources as a variable source, without considering the host behavior. This approach has proved very useful to deal with the so-called retroactivity [21], the loading interaction among circuit modules and host originated from mass exchange. Retroactivity poses problems when predicting the behavior of a large network from that of the composing modules. It is a problem analogous to modeling the coupling between electrical circuits connected to a real energy source. Thus, the models accounting for it somewhat resemble Ohm’s law [6, 31]. As these models do not explicitly consider the host behavior they cannot be easily used within a multiscale framework integrating the synthetic circuits of interest, the host, and the cell environment at the macroscopic level.

Alternatively, one may develop models relating substrates uptake, cell growth rate, and availability of free resources as a function of the gene circuits demand. These range from very coarse-grain ones [35, 16, 4] to semi-mechanistic ones with varied degrees of granularity [43, 18, 26]. In this last case, the interplay between circuit, host and environment can be directly incorporated into the circuit model of interest to capture the impact of cellular trade-offs and resource competition on the circuit function.

Construction of a large-scale mechanistic model of *E. coli* in [26] enabled the authors to integrate and cross-evaluate a massive, heterogeneous dataset integrating measurements reported by various groups over decades. On the other hand, medium-size detailed mechanistic models like the one developed in [43] have been used to study behavioral modulations of a gene switch [2] or a feedforward circuit [3, 15]. These medium and large-scale models, though very useful, are most often over-parametrized and cannot easily be integrated within a user-friendly lite and agile computational framework for model-based circuit design.

The main goal of the small-size model presented here is to provide enough granularity so as the model to provide good predictions of the dynamics of the host cell, the expression of the genes of interest and their interactions while having a small number of differential equations and parameters. An additional goal is to provide a model amenable for model-based circuit design purposes. To this end, the model considers explicitly lab-accessible gene characteristics such as promoter and RBS strengths and degradation rates of mRNA and proteins.

The rest of the paper is organized as follows. In Section 2 we derive the expression for the dynamics of protein expression considering the process of parallel translation of the same transcript by several ribosomes the resources recruitment strength functional coefficient. In Sections 3 and 4 we derive the dynamics of the number of mature available ribosomes and its relationship with the number of free ribosomes. This allows to derive the dynamics of protein expression as a function of the unbound fraction of available mature ribosomes In Section 5. These results are used in section 7 to derive an expression of the cell growth rate as a function of the number of ribosomes actively translating. The relationships between cell and population growth rates and between growth rate and cell mass are considered in sections 7 and 8 respectively. This first part of the paper finishes with the derivation of the equivalence between the relative resources recruitment strength and the relative mass fraction of a given protein at steady state, the average host dynamics and steady state and the evaluation of protein mass productivity in sections 9, 10 and 11 respectively. Next, we use experimental data from the literature and apply the previous developments to *E. coli*. We estimate the fraction of ribosomes being used in translating complexes relative to the mature available ones in section 13 and the average resources recruitment strength for both ribosomal and non-ribosomal *E. coli* proteins in section 14. These allow to estimate the number of free ribosomes that explain the experimental translation efficiencies per mRNA in section 15. In the last part of the paper we first show in sections 16 and 17 how the sensitivity of the resources recruitment strength to RBS and promoter can explain the variation of the cell mass distribution with growth rate and the differential roles of RBS and promoter strengths. Finally, in section 18 we show how host-circuit interaction shapes the optimal productivity of both endogenous and exogenous proteins in the expression space and we draw some brief conclusions in the last section.

## 2. Modelling protein expression

In our model, we consider a set of basic assumptions:

1. Transcription dynamics is fast enough as compared to translation so it can be considered at quasi-steady state (QSS).
2. The main resources-dependent process in protein expression is translation. Therefore:
  a. only ribosomes are considered as limiting shared resource required for protein expression. RNA polymerase is not considered explicitly.
  b. the effective translation rate is assumed to depend on the availability of intracellular substrate. This is implicitly considering that building the polypeptide protein chain is the limiting energy-consuming process in the cell. We do not explicitly consider the catabolic conversion of substrate into aminoacid building blocks.
3. Several ribosomes may translate a single messenger RNA (mRNA) simultaneously.
4. We identify a transcriptional unit by its promoter.

With these assumptions in mind, we model the expression of a given protein *p_k_* by means of the set of pseudo-reactions 1.

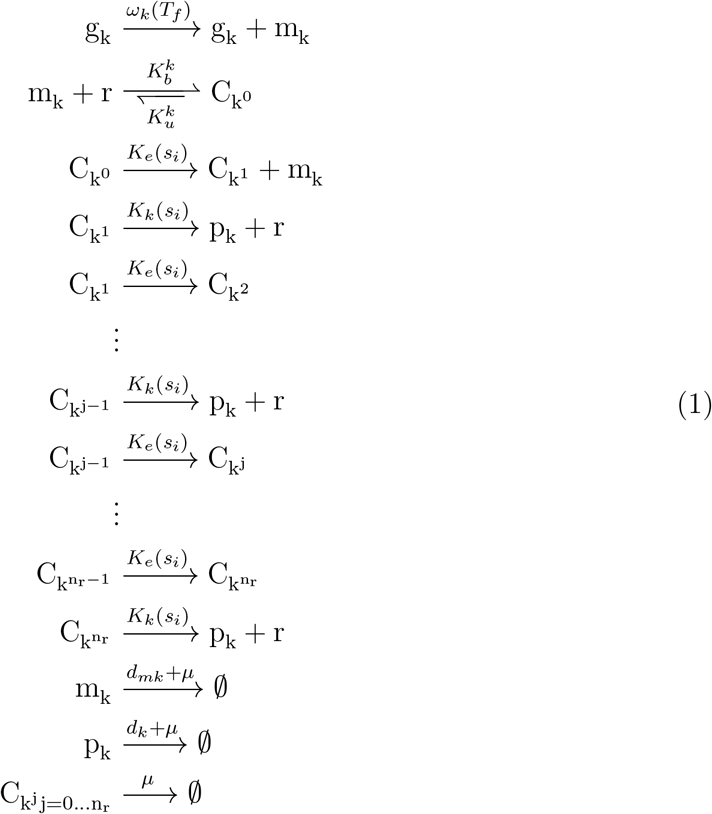

where the species involved are:

*g_k_* : free copy of the promoter, ie. of the gene transcriptional unit.
*m_k_* : free ribosome binding site (RBS), ie. mRNA copy with its RBS free.
*r* : free ribosome.
*C*_*k*^0^_ : complex formed by a ribosome bound to the RBS in a mRNA
*C_k^j^_* : *j*-th translating complex formed by *j* ribosomes simultaneously translating a mRNA copy with freed RBS.
*p_k_* : protein copy number
*s_i_* : intracellular substrate (molecules)

and the specific reaction rates stand for:

*ω_k_*(*T_f_*) : transcription rate. It may be a function of one or several transcription factors (TF) and may include the gene copy number.
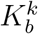 : association rate between a free ribosome and the RBS.
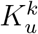 : dissociation rate between a free ribosome and the RBS.
*K_e_*(*s_i_*) : translation initiation rate.
*K_k_*(*s_i_*) : translation elongation rate.
*d_mk_* : mRNA degradation rate.
*d_k_* : protein degradation rate.
*μ* : cell specific growth rate.

For a protein of length *l_pk_* aminoacids, we denote:

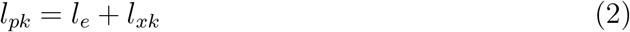

where 1/*l_e_* is the ribosomes density, with *l_e_* expressed as equivalent number of codons. We consider the effective RBS length to be the same as *l_e_*. The remaining length of the protein is denoted as *l_xk_*. Thus, up to *n_r_* ribosomes can simultaneously be translating a single copy of mRNA, with *n_r_* = *l_xk_*/*l_e_*, and an additional ribosome is bound to the RBS.

We consider the effective rates *K_e_*(*s_i_*) and *K_k_*(*s_i_*) at which the ribosome glides through the RBS and the remaining mRNA nucleotides respectively. Thus, we consider:

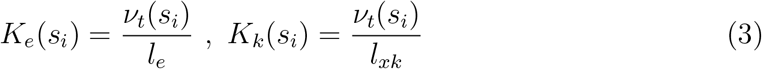

with

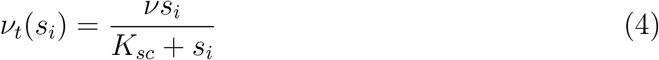

where *v* is the maximum translation rate and *K_sc_* is a Michaelis-Menten parameter related to the cell substrate uptake and catabolic capacity. As a first approximation, we consider that *ν* is organism dependent but does not depend on the sequence of nucleotides, and *K_sc_* is organism and substrate dependent but does not depend on the nucleotides sequence either.

The translating complexes *C*_*k*^*j*–1^_ are pseudo-species modeling the process of parallel translation. They can loosely be identified with each of the chains of aminoacids under formation. The first one, *C*_*k*^0^_, represents the ribosome bound to the RBS. Notice with rate *K_e_*(*s_i_*) the ribosome bound to the RBS, forming *C*_*k*^0^_, advances to the next ribosome occupancy slot, generating the translating complex *C*_*k*^1^_ and freeing the RBS so a new ribosome can enter in the cue. In turn, the displacement of the ribosomes in the cue by one occupancy generates the translating complexes *C_k^j^_* from the previous complex *C*_*k*^*j*–1^_. Finally, each parallel translating complex *C_k^j^_* generates a protein with rate *K_k_*(*s_i_*) and frees its bound ribosome. Recall the translating complexes can be identified with each of the chains of aminoacids under formation. This way, we decouple the cue dynamics of the ribosomes advancing along the mRNA from the protein building ones, thus getting a continuous approximation of the parallel process of translation.

Next, we apply mass action kinetics to obtain the dynamic balances for the copy number of each species in the model. This way, we have:

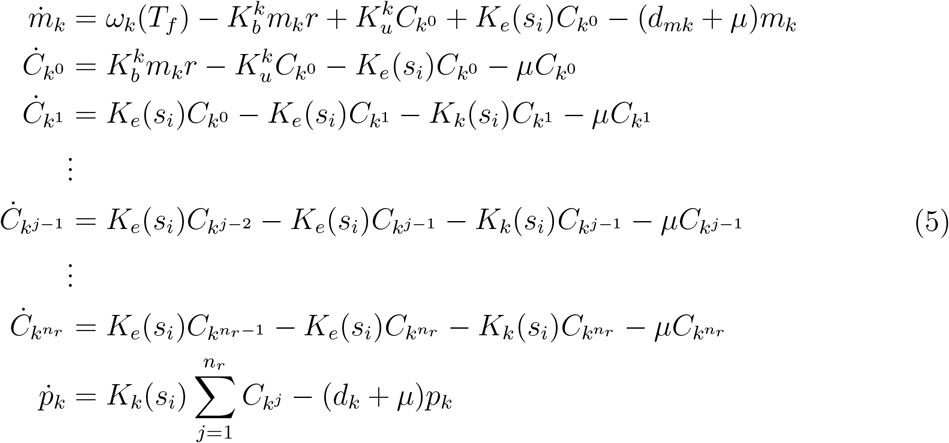

Recall we assume that transcription dynamics is fast enough as compared to translation so it can be considered at quasi-steady state (QSS). We also assume the binding-unbinding dynamics to form the translation complexes *C_k^j^_* are fast enough so that we can also consider the copy number of each of the complexes quickly reaches steady state. Therefore, from 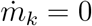 and 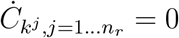 we get:

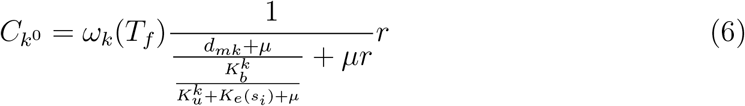

and

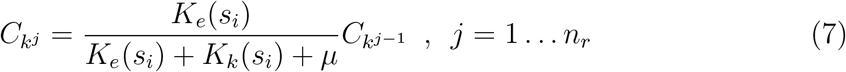

In practice, *d_mk_* ≫ *μ* and 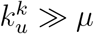 (see Table 12). Therefore the magnitude of the specific growth rate *μ* can be neglected with respect to both the mRNA degradation rate *d_mk_* and the sums 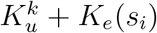 and *K_e_*(*s_i_*) + *K_k_*(*s_i_*) respectively so that we can approximate:

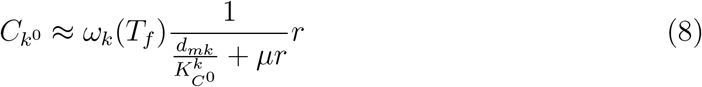

and

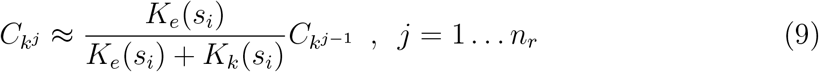

where we have defined:

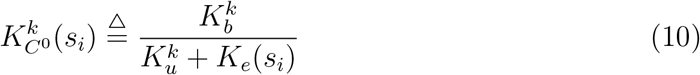

Notice 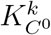 is directly related to the RBS strength.

Now, the dynamics for the copy number of the protein *p_k_* can be obtained from the equations (5) and (9) as:

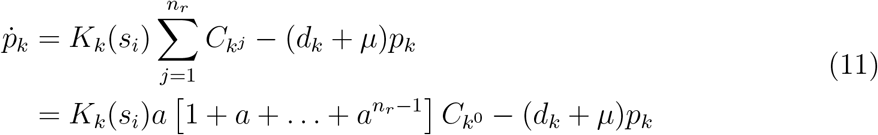

with 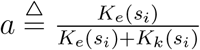.

Notice the geometric sum 1 + *a* + … + *a*^*n_r_*–1^ converges, as *a* < 1, so that:

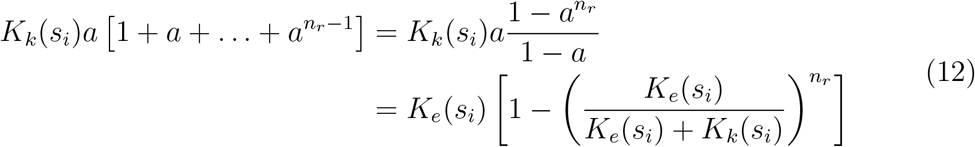

Using the definitions (3), notice 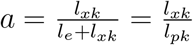 and we get the expression:

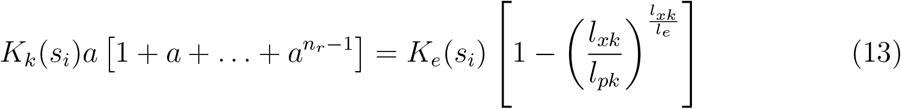

where we have taken into account that the maximum number of ribosomes bound to active translating complexes 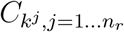 simultaneously translating a mRNA molecule is *n_r_* = *l_xk_*/*l_e_*, and we assume this maximum is always reached.

Recall we have assumed that transcription is not a limiting process, so we can express the effective transcription rate:

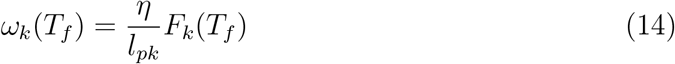

where *η* (aa/min) is the maximum transcription speed and *F_k_*(*T_f_*) is the transcription characteristic function that may depend on one or several transcription factors. By default, we assume the gene copy number *c_nk_* is one. If this is not the case, the effective transcription rate *ω_k_*(*T_f_*) must be multiplied by *c_nk_*. Notice in this case the transcription characteristic function *F_k_*(*T_f_*) may depend on the gene copy number as described in [41].

As commented in our preliminary assumptions, we do not model competition for RNA polymerases, which would also affect the effective transcription rate. Yet, notice that, even if there are no cognate transcription factors associated, the term *F_k_*(*T_f_*) can be used to account for competition for RNA polymerases preventing the effective transcription from proceeding at its maximum rate 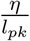, sequence-dependent affinity of the promoter for the RNA polymerases (promoter strength) and the effect of nucleotides usage on the transcription speed. In summary, the term *F_k_*(*T_f_*) can be used to accommodate aspects affecting transcription so that *ω_k_*(*T_f_*) is the effective transcription rate.

Thus, the dynamics for protein expression become:

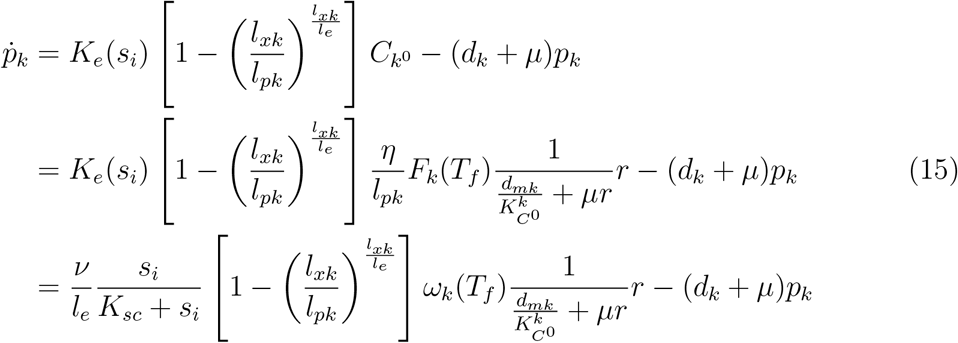

We now define the risosomes density related term:

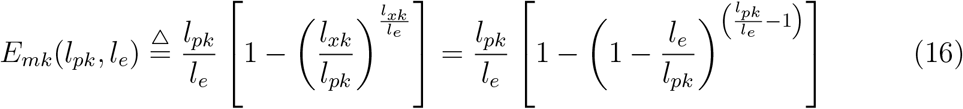

Figure 1 shows the values of *E_mk_*(*l_pk_, l_e_*) as a function of the protein length *l_pk_* for different values of *l_e_*. As, seen *E_mk_*(*l_pk_, l_e_*) can be accurately approximated as a linear function of *l_pk_*:

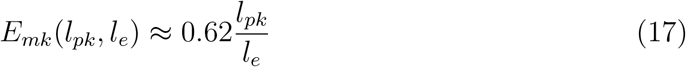

**Figure 1:**
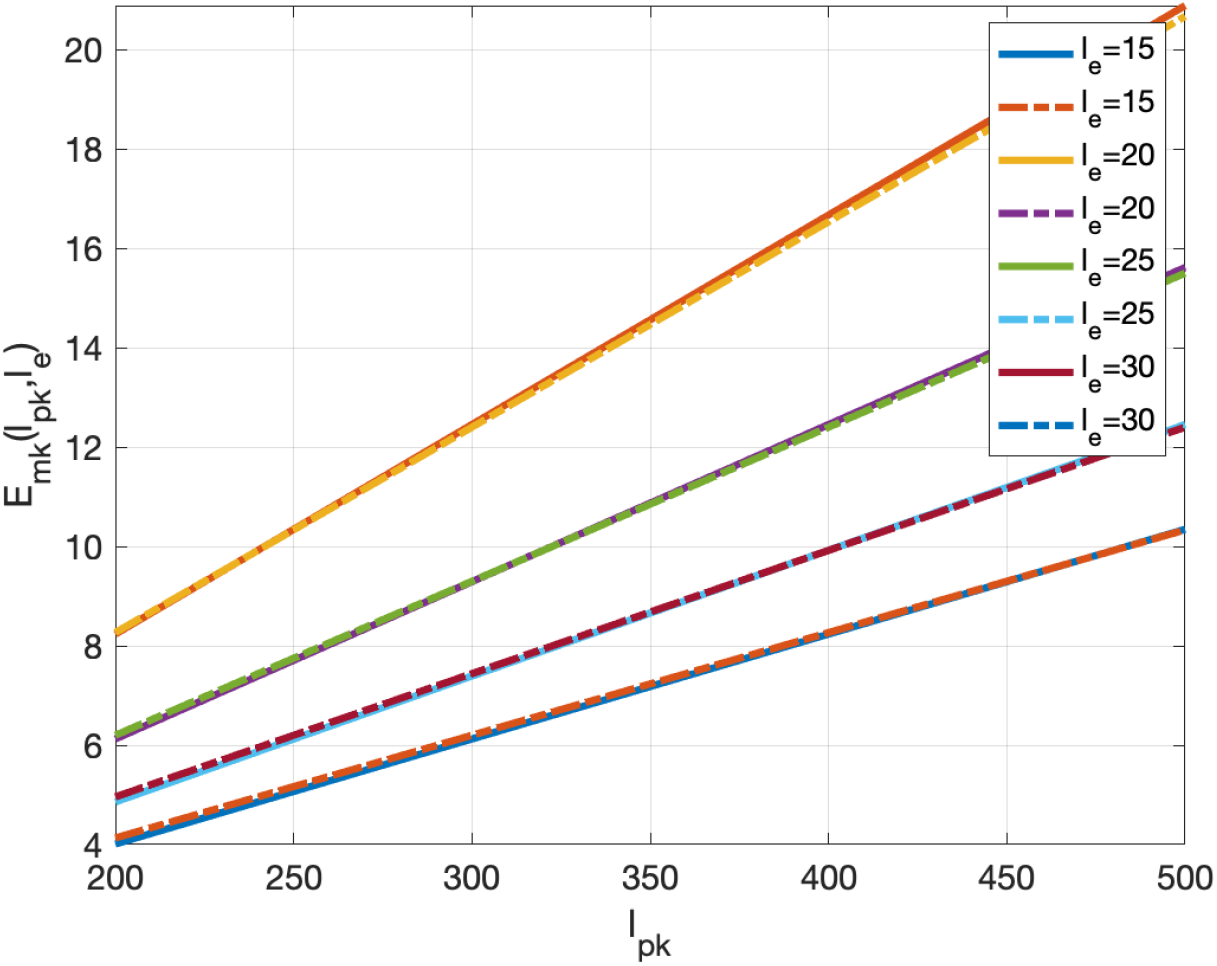
Function *E_mk_*(*l_pk_, l_e_*) as a function of the protein length *l_pk_* for different values of *l_e_* and their corresponding linear approximations 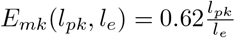.

We further define the *resources recruitment strength* functional parameter *J_k_*(*μ, r*):

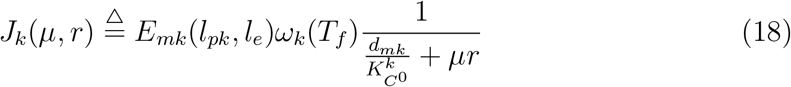

Notice the *resources recruitment strength* is a dimensionless function that expresses the capacity of the *k*–th gene to recruit cellular resources to get expressed. It explicitly depends on:

- the gene characteristics:
  – mRNA transcription and degradation rates *ω_k_*(*T_f_*) and *d_mk_*
  – RBS strength-related parameter 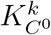
- and the availability of cell resources:
  – flux of free ribosomes *μr*
  – ribosomes density 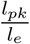 (via *E_mk_*(*l_pk_, l_e_*))

Using these definitions in equation (15) we get the expressions for the copy number dynamics of the *k*–th protein:

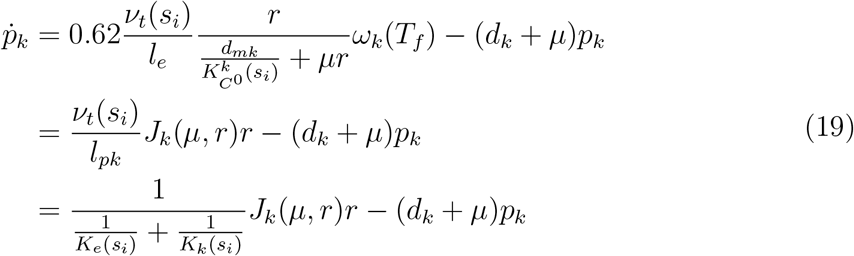

### Note

Using an electrical analogy, notice the term 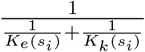 corresponds to the equivalent resistance of the circuit formed by the parallel resistances *K_e_*(*s_i_*) and *K_k_* (*s_i_*).

## 3. Dynamics of the number of mature available ribosomes

Ribosomes are large complexes formed by both ribosomal RNA molecules and a variety of ribosomal proteins, adding up to 55 proteins in *E. coli*. Recall we consider that translation is the main energy and resources limiting process we model. Notice the total copy number of ribosomes in the cell at any one time instant *r_t_* is the sum of the mature (*r_a_*) and inmature (*r_i_*) ribosomes. In turn, the mature ribosomes *r_a_* available for protein translation comprise the free ribosomes *r* and the ones already bound to translating complexes being used to build proteins. These comprise both the ribosomes *r_r_* bound to translating complexes building the ribosomes themselves, and the ones *r_p_* bound to the translating complexes of non-ribosomal proteins. These last will comprise those associated to the gene circuits of interest (*r*_sys_), and the ones used by the rest of the transcriptional units being expressed in the cell. Under nominal conditions, these last represent the basal needs of the cell and hereafter we will refer to them as the wild-type (wt) bound amount of ribosomes (*r*_wt_). Thus, we have *r_a_* = *r* + *r*_bound_, with *r*_bound_ = *r_r_* + *r_p_* and *r_p_* = *r*_sys_ + *r*_wt_.

The copy number of available mature ribosomes is a fraction of the total number of ribosomes, so that *r_a_* = Φ*_t_r_t_*. The fraction Φ_*t*_ varies little in time, with an average value Φ_*t*_ = 0.8 [4, 5] so that the dynamics of the total number of ribosomes and that of the number of available ribosomes are the same but for a scale factor. Next we consider the dynamics of the total copy number of ribosomes in the cell, *r_t_*.

To get the dynamics of *r_t_* we first consider an analogous expression to (19) for each of the proteins forming up a ribosome. If we consider the average ribosomal protein *p_r_*, the total number of ribosomal proteins for a given average ribosome composed of *N_r_* proteins (e.g. *N_r_* = 55 for *E. coli*) is defined as *p_Σr_* = *N_r_p_r_*. For the average ribosomal protein *p_r_* we have the dynamics:

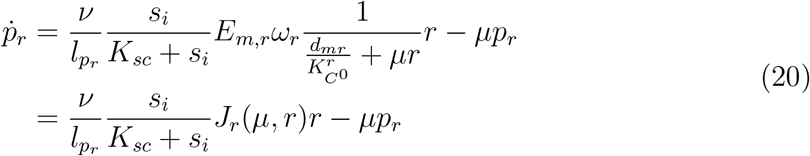

where we have assumed that ribosomal proteins are only subject to dilution caused by cell growth.

Then, the dynamics for the total number of ribosomal proteins can be approximated as:

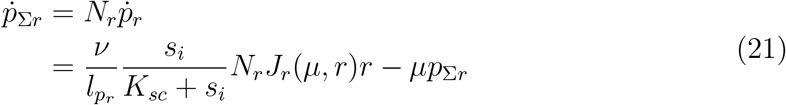

Since we need all *N_r_* proteins forming up an individual ribosome, and considering the average ribosomal protein *p_r_*, the total copy number of ribosomes is *r_t_* = *p_Σr_/N_r_*. Therefore, the dynamics of the total copy number of ribosomes *r_t_* will be the same as those of *p_r_*. That is:

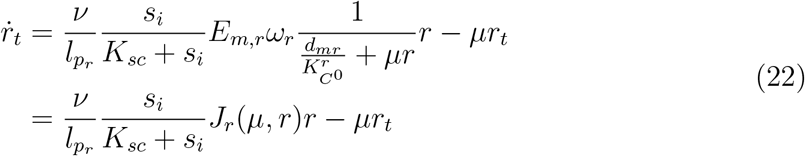

Therefore, the dynamics of the number of mature available ribosomes is:

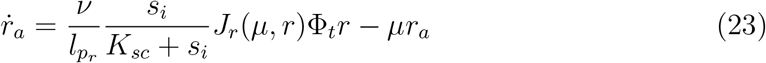

## 4. Relating free and available ribosomes

Recall we had *r_a_* = *r* + *r*_bound_. For each protein *p_k_* of interest, and using the previous results, the number of ribosomes bound to complexes involved in its translation at each time instant is given by:

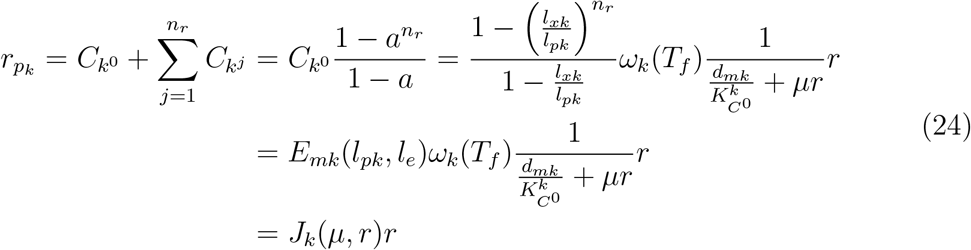

An analogous expression can be obtained for the number *r_r_* of ribosomes bound to complexes involved in translation of ribosomes themselves:

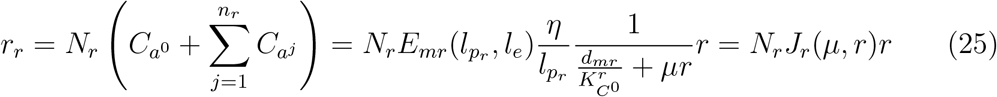

where we have taken into account that it requires *N_r_* proteins to build-up a ribosome.

Therefore, the copy number of mature available ribosomes *r_a_* can be obtained from:

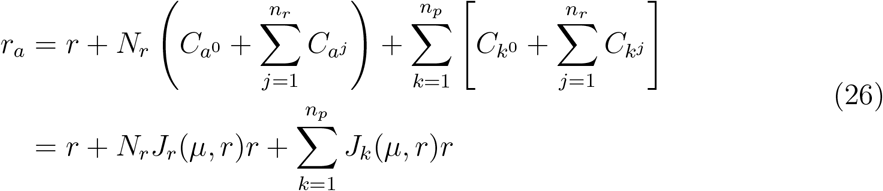

where *n_p_* = *n*_sys_+*n*_wt_ is the sum of the quantities of basal wild-type and system-of-interest non-ribosomal proteins in the cell.

From equation (26) we get the number of free ribosomes *r* as a function of the mature available ones *r_a_*:

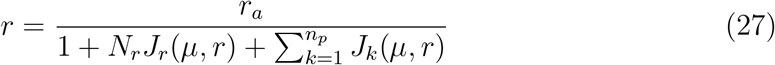

## 5. Protein expression dynamics as a function of the unbound fraction of available mature ribosomes

We can define:

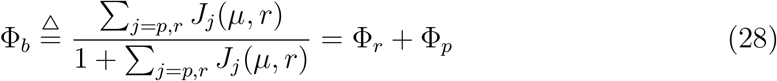

with

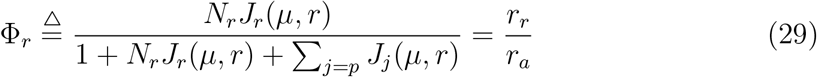

and

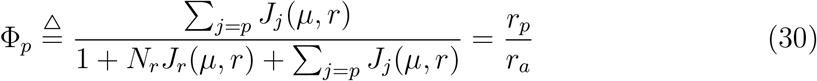

where we have used (24), (25) and (27) and 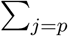 stands for the sum over the ensemble of all non-ribosomal proteins *p* = 1…*n_p_*.

Notice that

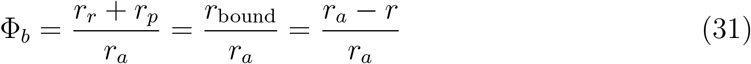

is the fraction of ribosomes bound to translating complexes (including both translating complexes for ribosomal and for non-ribosomal proteins) relative to the available mature ribosomes.

Notice each term

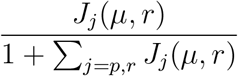

in Φ_*p*_ can be understood as the share of cell resources required by the *j*–th protein. Thus, the magnitude of the adimensional coefficient *J_j_*(*μ, r*) is a measure of the resources recruited by the *j*–th protein.

Using the definitions (14), (16), (18) and (28) and taking into account the relationships above, we can express the dynamics of protein expression(19) and available ribosomes (22) as:

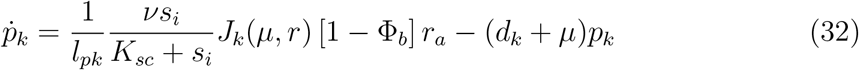

and

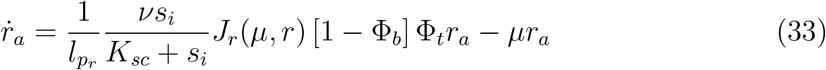

respectively.

As for the dynamics of the total number of ribosomes, using *r_a_* = Φ_*t*_*r_t_* we have:

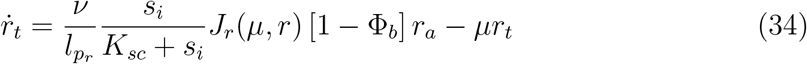

## 6. Obtaining the cell specific growth rate

Cell growth can essentially be explained as the time variation of the protein fraction of the total cell mass. To deal with protein degradation, we take into account that the protein fraction of cell mass is the sum of the mass of functional and non-functional proteins (ie. proteins undergoing degradation). Though non-functional proteins do not contribute to cell growth, they do to cell mass. Thus, for a protein *k* we can consider the fraction quantity of functional molecules of the protein, *p_k_*, and the one of non-functional units 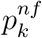 so that the total number of proteins of the species *k* is 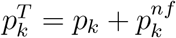. Then, considering the dynamics (32) of a generic protein, we will have:

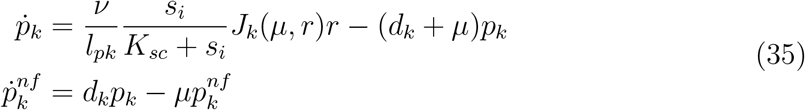

where we have taken into account that the non-functional fraction 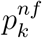 only undergoes dilution due to cell growth, there is a conversion from functional to non-functional fraction caused by protein degradation, and notice that all expressions for protein expression obtained in the previous sections referred to the functional fraction of proteins.

If we consider the average mass of an aminoacid *m_aa_*, the mass weight of a protein of length *l_pk_* can be approximated as *m_aa_l_pk_*. Thus, for 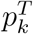 molecules of the *k*-th protein, their total mass weight is 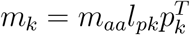.

The mass of *n_p_* proteins can be approximated as:

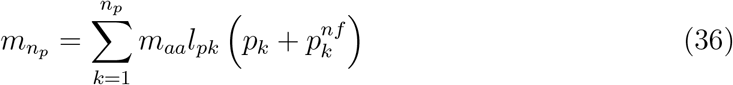

Therefore, the times variation of the protein mass explained by this set of *n_p_* proteins is:

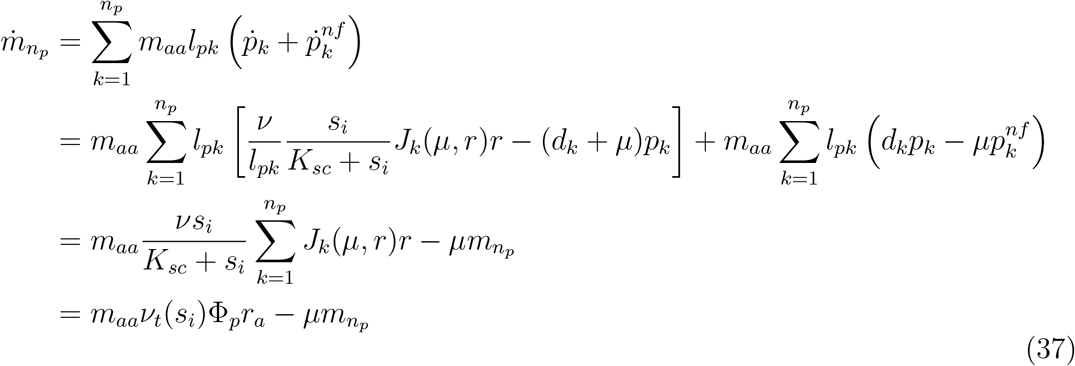

where recall *J_k_*(*μ, r*)*r* is the number of ribosomes bound to complexes involved in the translation of each *k*–th protein and we have used the expression (27) relating the free and available ribosomes and definition (30). Notice, in addition, that degradation of proteins does not play a role.

As for the protein cell mass variation explained by the time variation in the total number of ribosomes *r_t_* we will have the analogous expression:

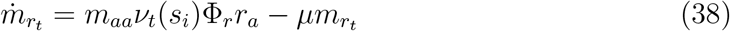

where we have used again the fact that *N_r_* ribosomal proteins are required to form up one ribosome. Notice that here we are only considering the weight of the protein fraction of the ribosomes. This accounts only for approximately one third of the ribosomes mass [28].

Denoting the cell protein weight *m_n_p_+r_t__* = *m_n_p__* + *m_r_t__*, we reach the expression:

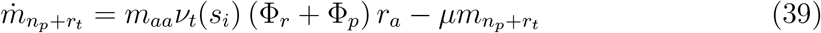

Now, consider the set *n_p_* of proteins being simultaneously expressed equals the sum of the proteins being expressed by the gene circuits of interest *n*_sys_ plus all remaining cell environment (or context) proteins expressed in the cell *n*_wt_. Then, the dynamics of the cell protein mass (*m_p_*) can be approximated as:

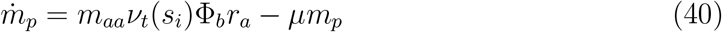

Recall that the specific growth rate *μ* is a continuous approximation of the discrete event process of cell duplication. Here we consider that the total biomass dry weight variation (ie. that of the whole population of cells) is mainly caused by cell duplication (i.e. population growth), and the dynamics of cell mass accumulation are much faster than those of cell duplication. Under this assumption, we may consider the protein mass for each cell quickly reaches steady state 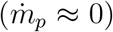. Thus, from equation (40) we get the expression for the cell specific growth rate:

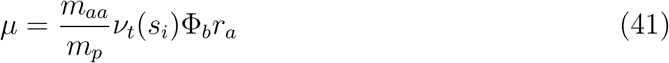

Notice Φ*_b_r_a_* is the number of ribosomes actively translating proteins (both ribosomal and non-ribosomal) at a given time instant. Equation (41) allows to predict this number given a specific growth rate, assuming saturation of intracellular substrate (eg. considering a batch experiment and the exponential growth phase) and considering the average values for the amino acids mass and that of the protein fraction of the cell.

Notice (41) can be expressed as a function of the total number of ribosomes *r_t_* as:

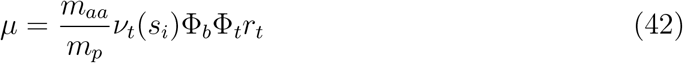

## 7. Cell specific growth rate and population dynamics

Now, it will be interesting to relate the expression for the cell specific growth rate *μ* obtained in (41) with the classical Monod-like expressions for a population of cells growing in a bioreactor.

Recall if we have a population of *N* cells and we consider the average cell dry mass *m_c_*, the total biomass dry-weight will be *M_p_* = *Nm_c_*. By taking derivative with respect to time, we get:

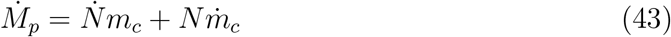

Now, consider the continuous approximation of cell duplication:

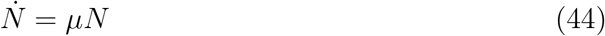

where *μ* = log2/*t_d_*, with *t_d_* the duplication time, is the specific growth rate. Then:

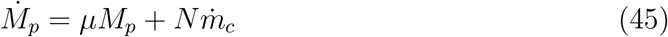

from which we get:

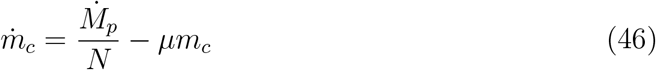

where 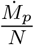 is the mean value per cell of the population mass growth.

As done in section 6, we assume the cell mass quickly reaches steady state as compared to the population dynamics. Thus, from equation (46) and assuming 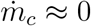 we get:

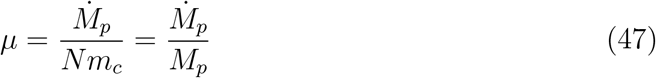

Experimental evidence suggests that the cell density *ρ* varies little throughout the adult cell life [20], so:

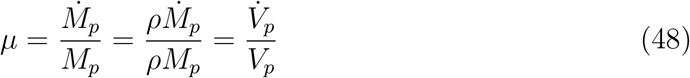

The identities (48) allow to relate the specific growth rate obtained in (41) with the one obtained from population-scale macroscopic experiments under the condition of steady-state growth where the rate of total cell-mass growth is identical to the rate of cell number growth [45]:

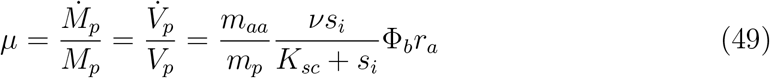

Next, we relate the expressions (49) for growth rate obtained from intracellular considerations with the classical empirical Monod expression for the specific growth rate under a limiting substrate obtained from the experimental analysis of a macroscopic culture:

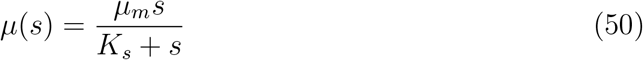

where *μ_m_* is the maximum specific growth rate, *K_s_* is the Monod affinity constant, and *s* is the concentration of the limiting substrate in the culture medium.

Alternative theoretical approaches to derive the Monod equation exist [24, 44, 42]. Here we follow a reasoning derived from the model developed in [43], where the quantity of intracellular substrate *s_i_* is related to the one of extracellular substrate *s* through the dynamics of nutrient import and catabolism:

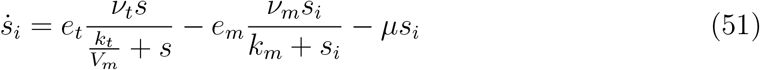

where *V_m_* is the volume of the culture broth, *e_t_* and *e_m_* are transport and catabolism enzymes, and Michaelis-Menten kinetics are assumed (see [43]).

If we assume that nutrient import quickly balances nutrient catabolism and we neglect the dilution term:

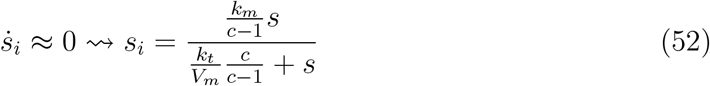

where 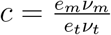. Notice if the maximum import and catabolism fluxes are balanced, ie. *c* ≈ 1, then there is a linear relationship between the intracellular amount of substrate and its extracellular concentration. Otherwise, if catabolism is more efficient than transport (*c* > 1) the intracellular amount of substrate *s_i_* saturates with increasing values of *s*.

Recall in our model the specific growth rate (49) is a function of *s_i_*. Using (52) we obtain the Monod-like expression:

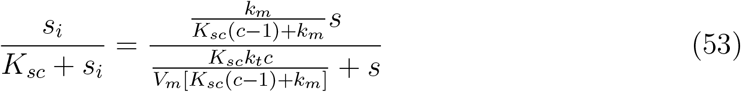

If we assume that the Michaelis-Menten constant for substrate catabolism is the same as the constant we defined in (3), that is, *K_sc_* = *k_m_*, we have:

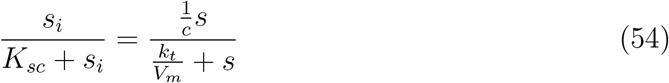

Notice that the hypothesis *K_sc_* = *k_m_* implicitly implies that the Michaelis-Menten constants for substrate catabolism and transport have similar values (*k_t_* = *k_m_*), in agreement with the assumptions in [43].

Also notice the term 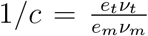 can be interpreted as the maximum flux yield between nutrient import and its catabolism. If *c* ≈ 1, that is, under the hypothesis that the efficiency of nutrient import and catabolism are balanced so the maximum import and catabolism fluxes are similar:

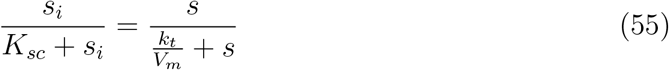

In case catabolism is more efficient than transport (*c* > 1) we will need an increase in the concentration of the substrate in the extracellular medium (*s*) to achieve the same value of 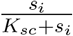 in (54) as compared to the balanced case *c* =1. Finally, in case transport is more efficient than catabolism (*c* < 1) we will need lower concentrations of the extracellular substrate.

### Note

The relationship (54) is valid but in the extreme cases non relevant cases *c* = 0 (there is transport into the cell but nutrients are not metabolised) and *c* = ∞. In these cases dilution cannot be neglected in equation (51) and the equilibria are different.

Then, from (49), (50) and (54) we get:

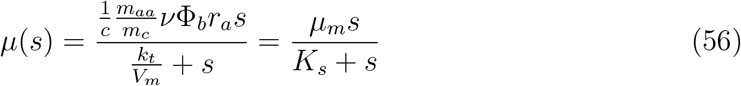

so we can identify:

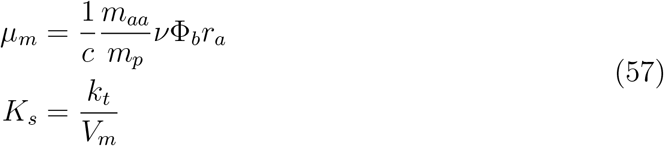

## 8. Relationship between growth rate and cell mass

Our model accounts for the protein mass distribution but does not consider the relationship between growth rate and the total cell protein mass. Several phenomenological models have been proposed in the literature accounting for the relationship between growth rate and cell dry weight, like the recent ones [37, 45]. In [45] a Monod-like relation between the chromosome replication–segregation period *C* + *D* and the cell specific growth rate *μ* is considered:

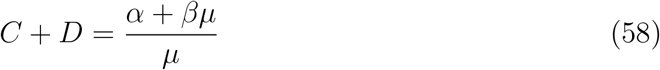

We estimated the parameters *α* and *β* in (58) using the data in [5]. Figure 2(left) shows the good fit obtained.

**Figure 2:**
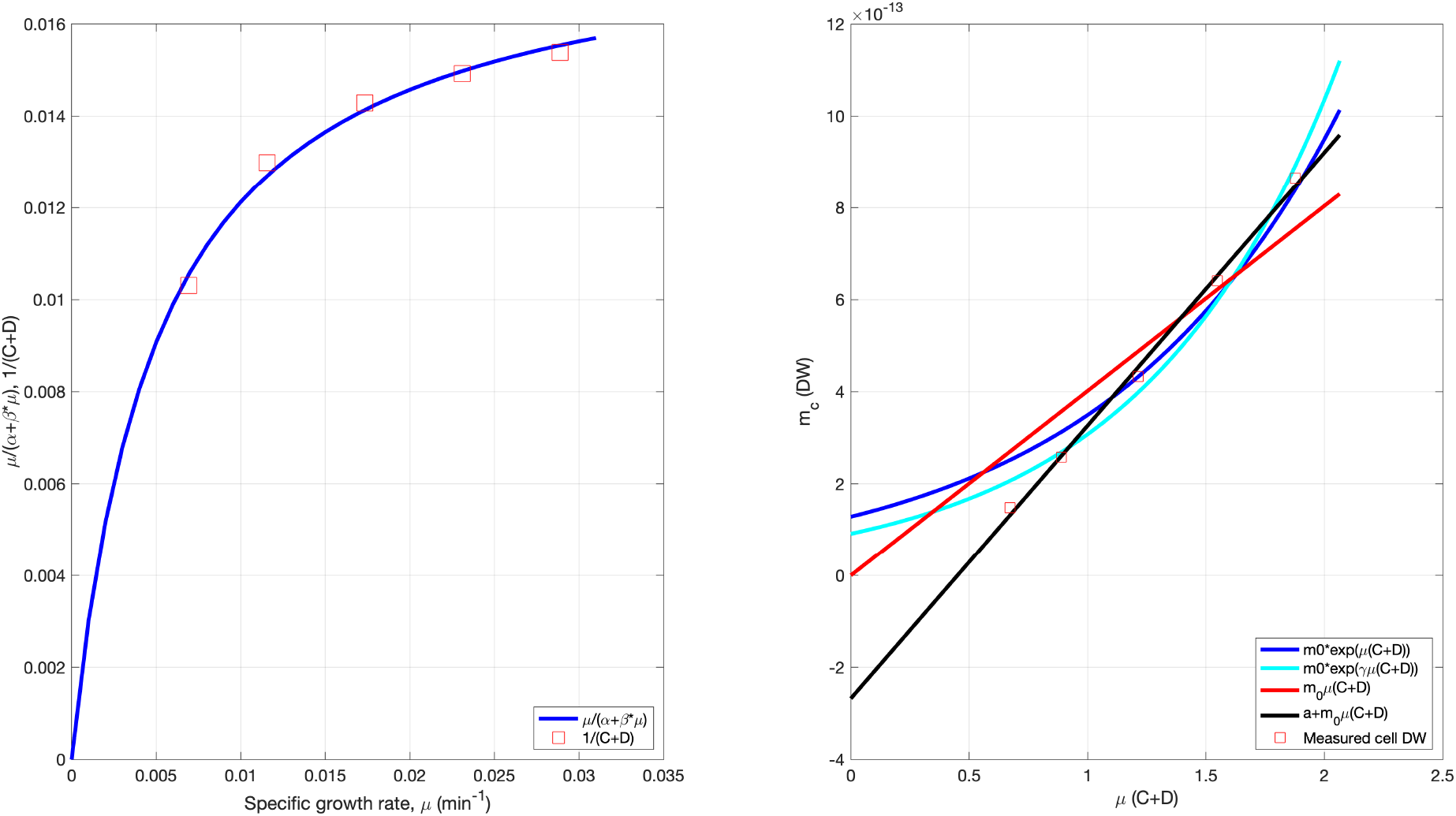
ns.

In addition the authors in [45] propose a linear relation between the cell dry weight and the product of *C + D* and *μ*:

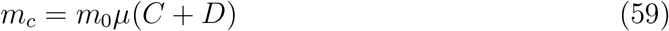

Therefore, according to this model, there is an affine relationship between the cell dry weight and specific growth rate:

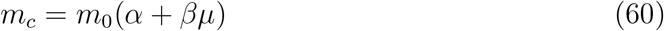

As shown in Figure 2(right), for the data we used, the relationship (59) gives a very rough approximation. Indeed, as shown in the same figure, and affine relationship gives much better fit.

As an alternative, in [37] an exponential relationship is proposed between the cell volume *S_c_* and the product of *μ*(*C* + *D*):

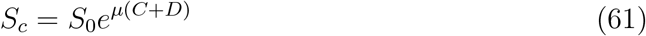

Assuming constant cell density, equation (61) can be expressed as a function of the cell dry weight. Figure 2(right) shows the fit assuming constant cell density, so that equation (61) can be expressed as a function of the cell dry weight. Notice that a better fit can be obtained considering the modified relationship:

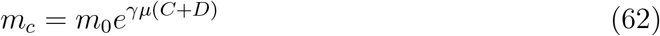

Now, if we consider expressions (58) and (62), we get:

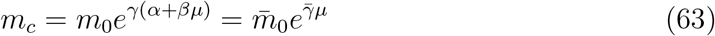

where 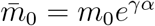 and 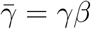.

Notice equation (63) is the solution of the differential equation:

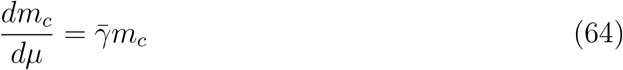

which expresses that the variation of cell mass (dry weight) relative to the variation of the specific growth rate is proportional to the cell mass. Figure 3 (left) shows the results obtained for (63) and Table 8 lists the best fitted parameters. Better fits can be obtained with alternative phenomenological expressions to (63) at the cost of losing the simple interpretation provided by expression (64).

**Figure 3:**
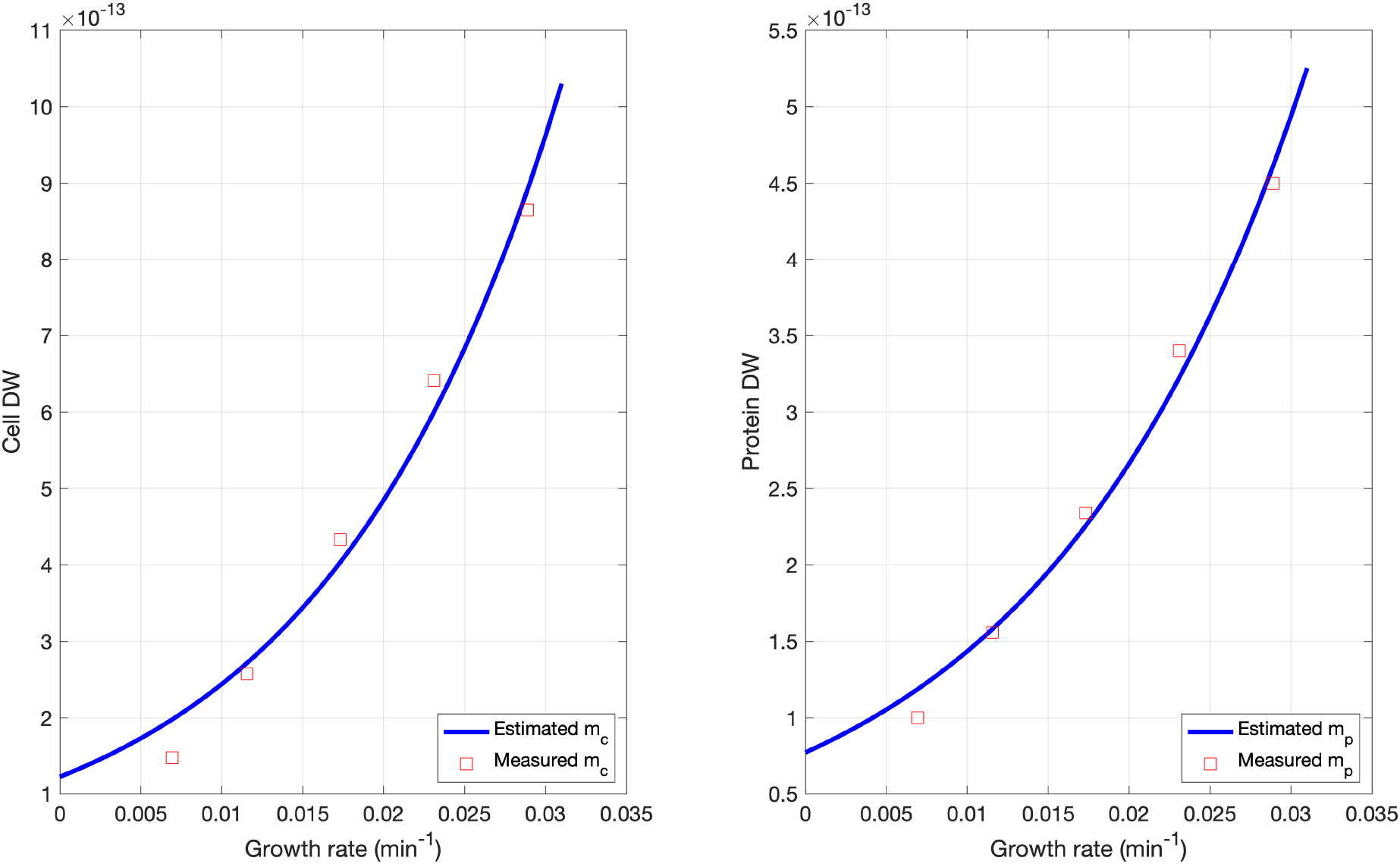
Fit between the cell dry weight (left) and protein content dry weight (right) and the specific growth rate.

For the relationship between the cell protein content and the specific growth rate we postulate a relationship analogous to (64). Thus, we consider:

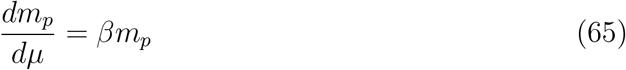

Figure 3 (right) shows the results obtained and Table 8 lists the parameters corresponding to the best fit.

Notice that alternative phenomenological relationships can be used instead, with possibly better fit to the experimental data. Thus, for instance, the power law model 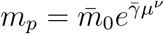 with 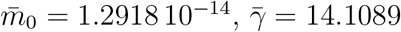 and *ν* = 0.389 is an alternative.

## 9. The relative resources recruitment strength provides the relative mass fraction at steady state

the relative resources recruitment strength of a given protein equals its relative mass

Recall the expressions (19) and (23) for the dynamics of given protein and the average ribosome respectively. Now, using (28), (41) and the dynamics of the total *k*–th protein 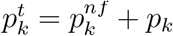 given by (35) we can rewrite:

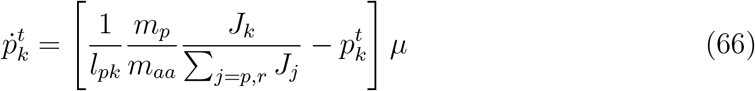

and

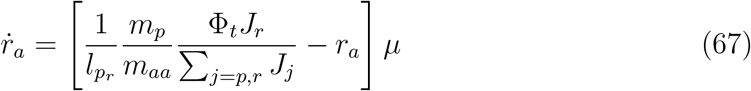

where 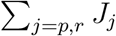 stands for the sum of the resources recruitment strengths for all proteins in the cell, including both ribosomal and non-ribosomal ones.

Now, using the expression for the cell protein mass

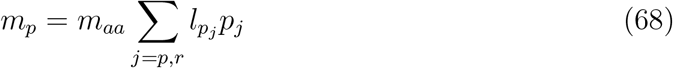

where the sum includes both the ribosomal and non-ribosomal proteins and we have assumed a common average amino acid mass *m_aa_*, we have:

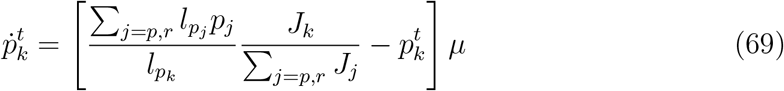

and

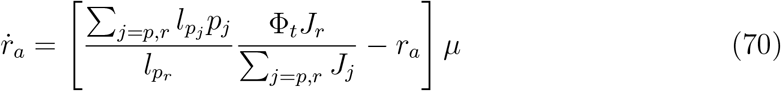

Notice that the steady state will reached either when the growth rate *μ* = 0 or at:

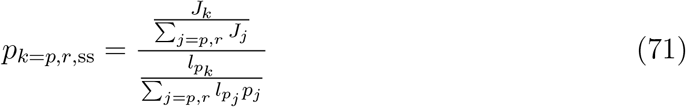

where *p*_*k*=*p,r*,ss_ stands for any protein, either ribosomal or non-ribosomal. Notice that expression (71) tells us that at steady state the relative resources recruitment strength of a given protein equals its relative mass in the cell:

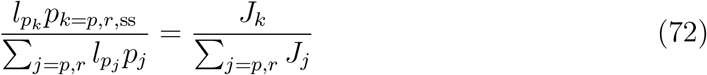

## 10. Average host dynamics and steady state

We were interested in having an average model for the host dynamics and its steady state that can be used as base for analysing host-circuit interactions. To this end we considered, on the one hand, the dynamics of the mass of total ribosomes *m_r_t__* and, on the other, the ones of the mass of the ensemble of non-ribosomal proteins *m*_wt_. In the later case, we have considered this set as a lumped species with a single average resources recruitment strength *J*_wt_(*μ, r*).

Thus, using (67), we have:

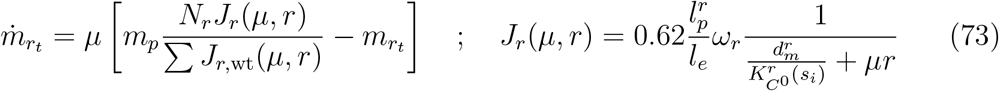

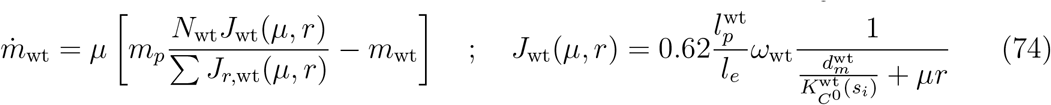

with the effective RBS strengths

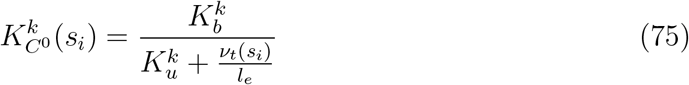

for *k* = {*r*, wt}, where we use the data of *ν_t_*(*μ*) for the effective translation rate *ν_t_*(*s_i_*)-implicitly assuming the growth rate at steady state is a function of the substrate *s_i_*– and where

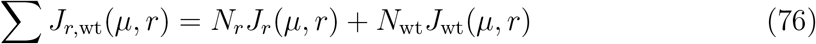

and *N_r_* and *N*_wt_ are the number of ribosomal proteins building up a ribosome (see Section 3) and the number of non-ribosomal proteins in the *wild type* cell respectively, so that the number of proteins and the resulting mass are related by:

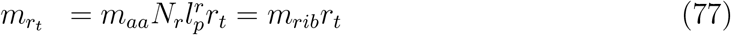

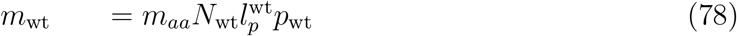

where *m_rib_* is the average mass of the protein content of ribosomes, and we have assumed a common average amino acid mass *m_aa_* as in (68) and average protein lengths 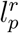 and 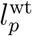.

Notice from (73)–(74) that at steady state we will have:

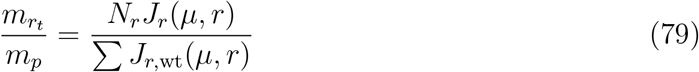

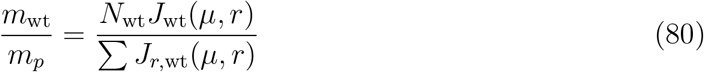

The number of free ribosomes *r* is obtained as:

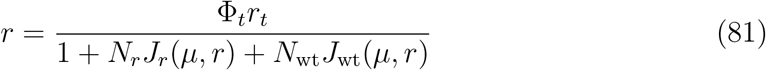

with *r_t_* obtained from (77), and the specific growth rate as:

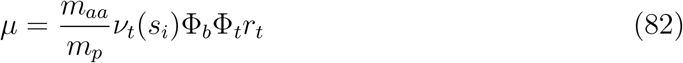

with

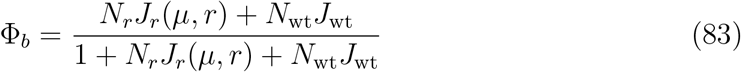

Notice from (73), (81) and (82) that at steady state:

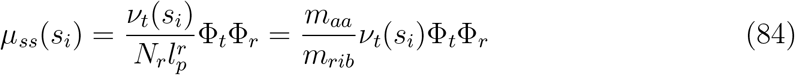

that is, the growth rate at steady state depends linearly on the fraction Φ_*t*_Φ_*r*_ of bound ribosomes being used to build up ribosomes relative to the total number of ribosomes.

At steady state, the flux of free resources for a given intracellular substrate *s_i_*, defined as the number of free ribosomes times the cell growth rate, can be obtained as:

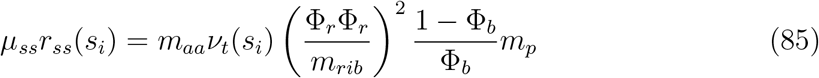

showing a linear relationship with the cell protein weight *m_p_*.

The cell protein weight *m_p_* depends of the growth rate. To this end, we used both the data available in [5] and the phenomenological relationships obtained in section 8.

## 11. Productivity of exogenous proteins depends on host-circuit interaction

Product titer and productivity are important measures of performance in biotechnological applications. In this section we consider the expression of a heterologous protein, and we analyse how its copy number at steady state and its specific productivity vary as a function of the RBS and promoter strengths and the host-circuit interaction.

We define the specific productivity a protein *A*, Π_*A*_, as the steady-state rate of expression of *A* per cell. Thus, if we consider a population of *N* cells and the continuous approximation of cell duplication (44), the total quantity of the protein *A*, 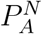, will increase as the population of cells does as:

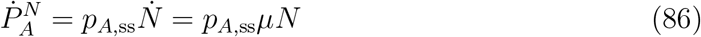

where *p*_*A*,ss_ is the quantity of the protein *A* at steady state in a single cell. Therefore:

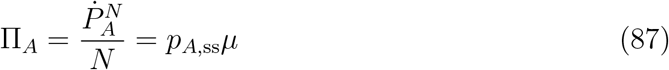

For the host endogenous dynamics we considered the model as described in section 10. We extend the model by adding the dynamics of the exogenous protein of interest A as:

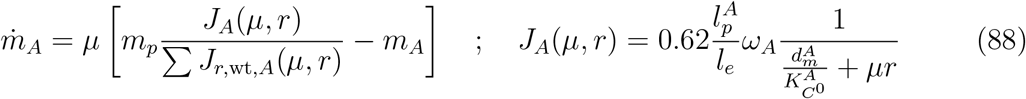

where

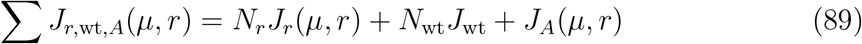

and the number of molecules of protein *A* and the resulting mass are related by:

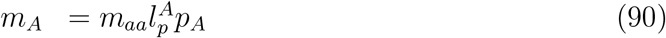

The number of free ribosomes *r* is obtained as:

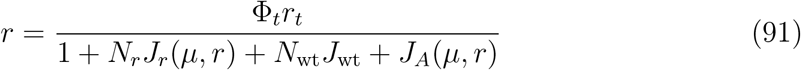

and the specific growth rate using (82) with

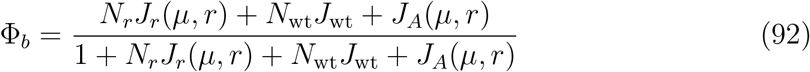

## 12. Model parameters

Table 12 shows the set of parameters used in the model.

## 13. Estimation of the fraction Φ*_b_*

Next we evaluate the fraction Φ_*b*_ of ribosomes being used in translating complexes relative to the mature available ones as a function of the free ribosomes *r* (see Section 5). We expect a very low number of free ribosomes. If this was not the case, there would be no real competition to recruit them. To set an initial upper limit, we use the estimation *r* = **N**(350, 35) in [23]. In addition, having too many free ribosomes in excess would imply a superfluous us of energy for the cell. Considering this hypothesis, we evaluated equation (31) as a function of the mature available ribosomes *r_a_* and the free ones r. Figure 4(left) shows the values estimated. Notice the values of Φ_*b*_ close to Φ_*b*_ = 1 indicating that the cell is always at the edge of its maximum capacity for using the available resources.

**Figure 4:**
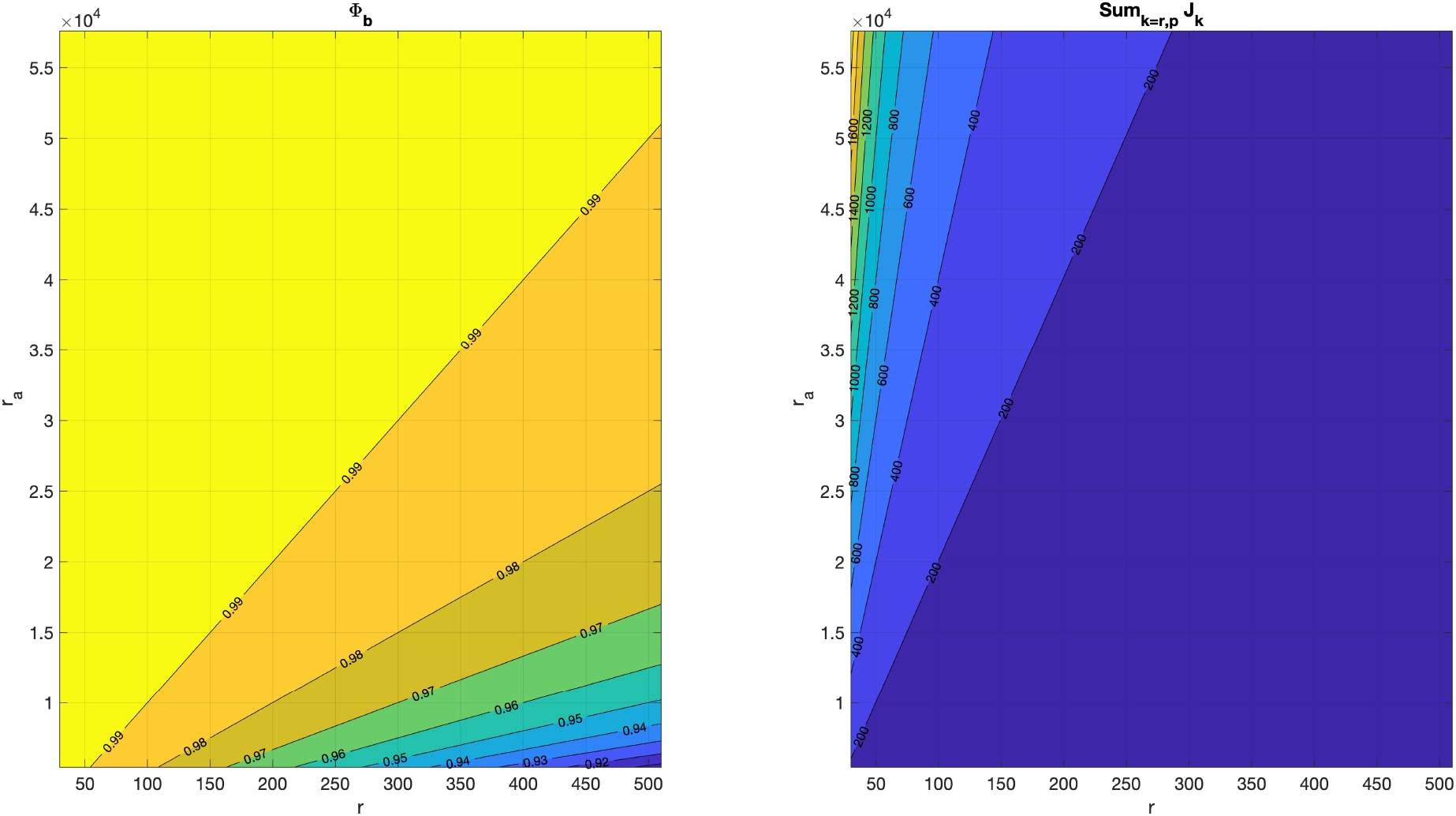
(Left:) Fraction Φ_*b*_ of ribosomes actually being used in translating complexes as a function of the available functional *r_a_* and the free ones *r*. (Right:) Corresponding estimation of the sum 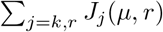

From the estimation of the fraction Φ_*b*_ we can obtain that of the dimensionless sum 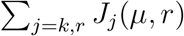 reflecting the resources recruitment load generated by the whole set of ribosomal and non-ribosomal proteins being expressed at a given moment in the cell (see Figure 4(right)) using the definition (28).

To validate the estimations above, we used the data in [19] to evaluate the sum 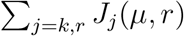 for all genes being expressed in *E. coli*. We took into account that the data in [19] was obtained for fast growing cells, with *t_d_* = 21 minutes. Under this conditions, it is sensible to consider that the intracellular substrate will be saturated and the cells are growing at its maximum growth rate. Then, we took into account that the number of active available mature ribosomes Φ_*b*_*r_a_* is related to the cell specific growth rate as described in Section 6. Recall, from equation (41) and considering substrate saturation, we have:

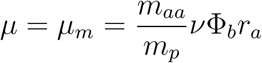

We estimated the cell protein mass *m_p_* as the total copy number of all proteins in the cell obtained in [19] times the average amino acid mass. The dynamic model for the expression of protein *p* in [19] considers:

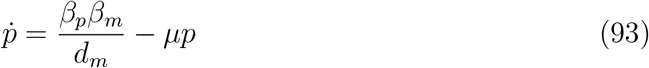

where *β_m_* (mRNA/t) is the transcription rate, *β_p_* (protein/(mRNA·t)) the translation one and *d_m_* the mRNA degradation rate. For those transcripts without information for *d_m_* in [19] we used the value shown in Table 12.

Then we used equation (19) to derive the relationship:

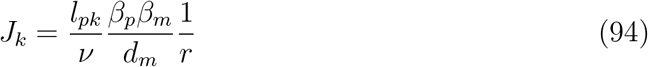

We evaluated (93) considering the set of all non-ribosomal proteins, the ribosomal ones, and the full set of proteins.

Figure 5(right) shows the very good agreement between the estimation obtained using equation (31) and the values calculated using equation (94) and the data in [19]. The difference is below 5.6% for all *r*. Is is important to stress that equation (31) is a purely theoretical expression. Only the expression (41) relating the number of active available mature ribosomes Φ_*b*_*r_a_* with the cell specific growth rate was used to estimate Φ_*b*_*r_a_* from the maximum specific growth rate corresponding to the data in [19].

**Figure 5:**
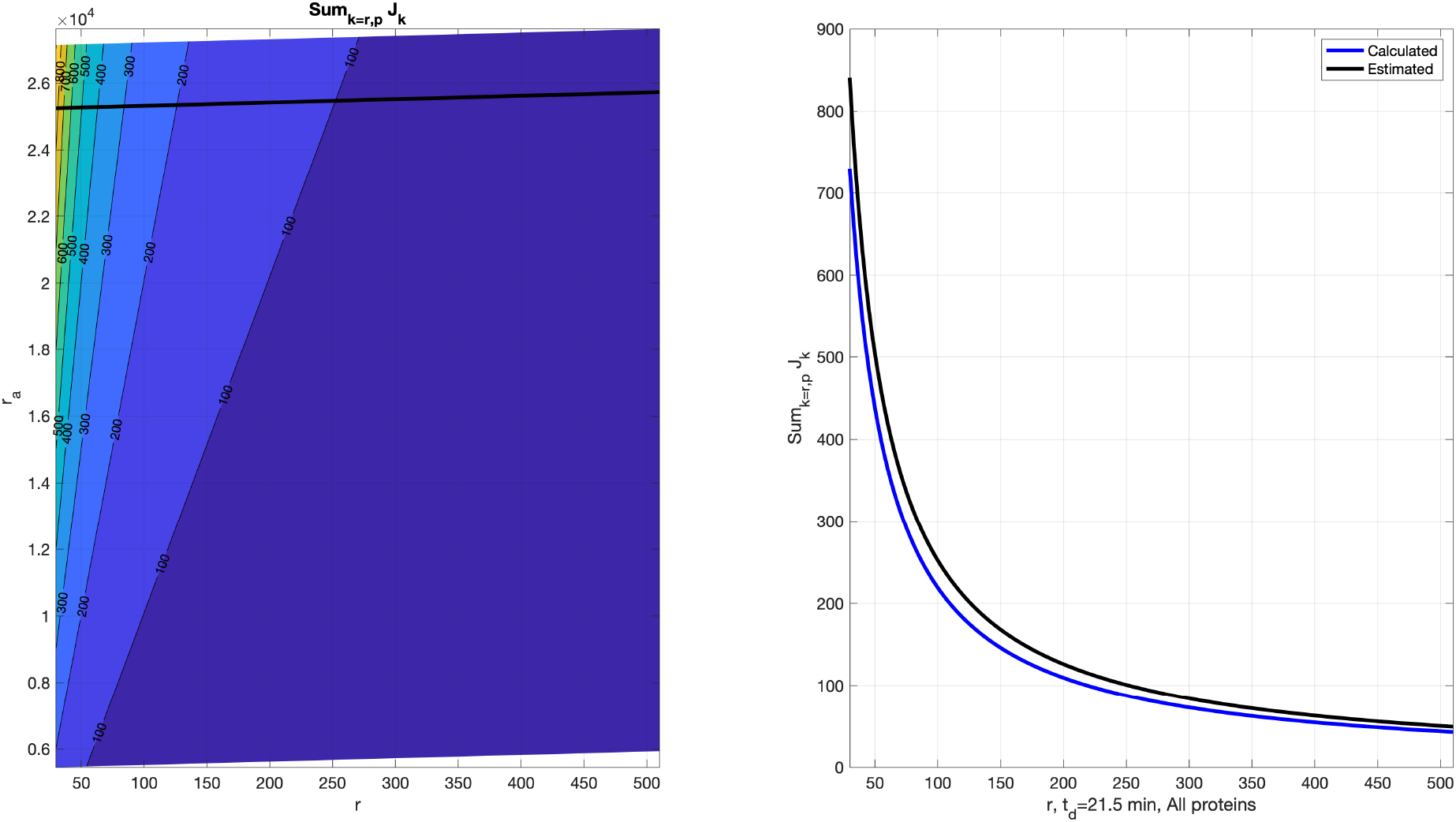
(Left:) Sum 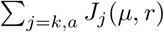 as a function of the available functional *r_a_* and the free ones *r*. The black line depicts the value of mature available ribosomes *r_a_* as a function of the free ones *r* assuming the number of bound ribosomes that correspond to the specific growth rate considered in [19]. (Right:) Corresponding estimation of the sum 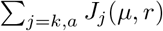 (black) and value obtained by evaluating the sum using (94) with the data in [19].

### NOTE

It is interesting to notice that from (41) and (31) we can derive the expression:

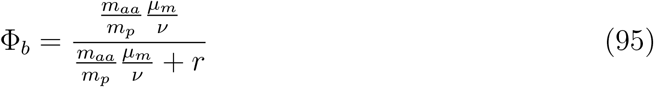

from which we obtain:

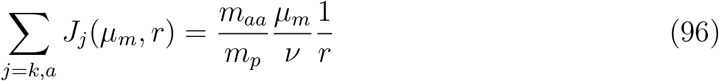

Now, considering (94), we get the relationships

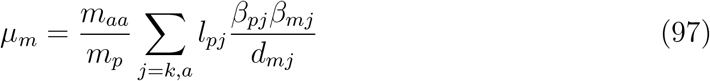

and

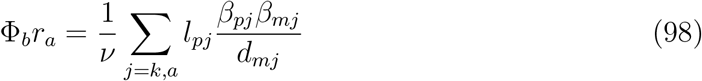

relating the maximum growth rate *μ_m_* and the bound ribosomes Φ_*b*_*r_a_* respectively with the transcription and translation rates, the transcripts degradation rates and the length of the proteins being expressed.

## 14. Estimation of the average resources recruitment strength

In Section 2 we derived equation (19) for the expression of a given protein and we defined the dimensionless function *J_k_*(*μ, r*) (see equation (18)) that quantifies the capacity of the *k*-th gene to recruit cellular resources to get expressed. In the previous section, we obtained the resources recruitment strength *J_k_* for each gene in *E. coli* estimated from the experimental data in [19], and the corresponding sum 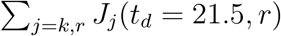 reflecting the total resources recruitment load in the cell. Next we evaluate the average resources recruitment strength *J*_avg_ as a function of the number of free ribosomes *r* for both ribosomal and non-ribosomal proteins.

As a first step we used the data in [19] and expression (94) to calculate the individual values of maximum resources recruitment strength *J_k_* evaluated at *r* =1 for the set of non-ribosomal and ribosomal protein expression genes and sorted them by magnitude. As seen in Figure 6 (top) the values of *J_k_* span several orders of magnitude. We also evaluated the ration between the sorted values of *J_k_* and the corresponding protein lengths. The results do not sensibly change with respect to the previous ones (see Figure 6 (bottom)). That is, the resources recruitment strength of *E. coli* genes is not fundamentally determined by the lengths of the proteins they code. This suggests, as expected, that factors such as the effective transcription and translation rates are more relevant. The fact that the ribosomal proteins, essential for the cell and continuously being expressed, have much higher resources recruitment strength values than the non-ribosomal ones. Moreover, the range of variation of *J_k_* over the ribosomal proteins is much lower than for non-ribosomal ones. This is consistent with the fact that to great extent all ribosomal proteins are equally important for the cell.

**Figure 6:**
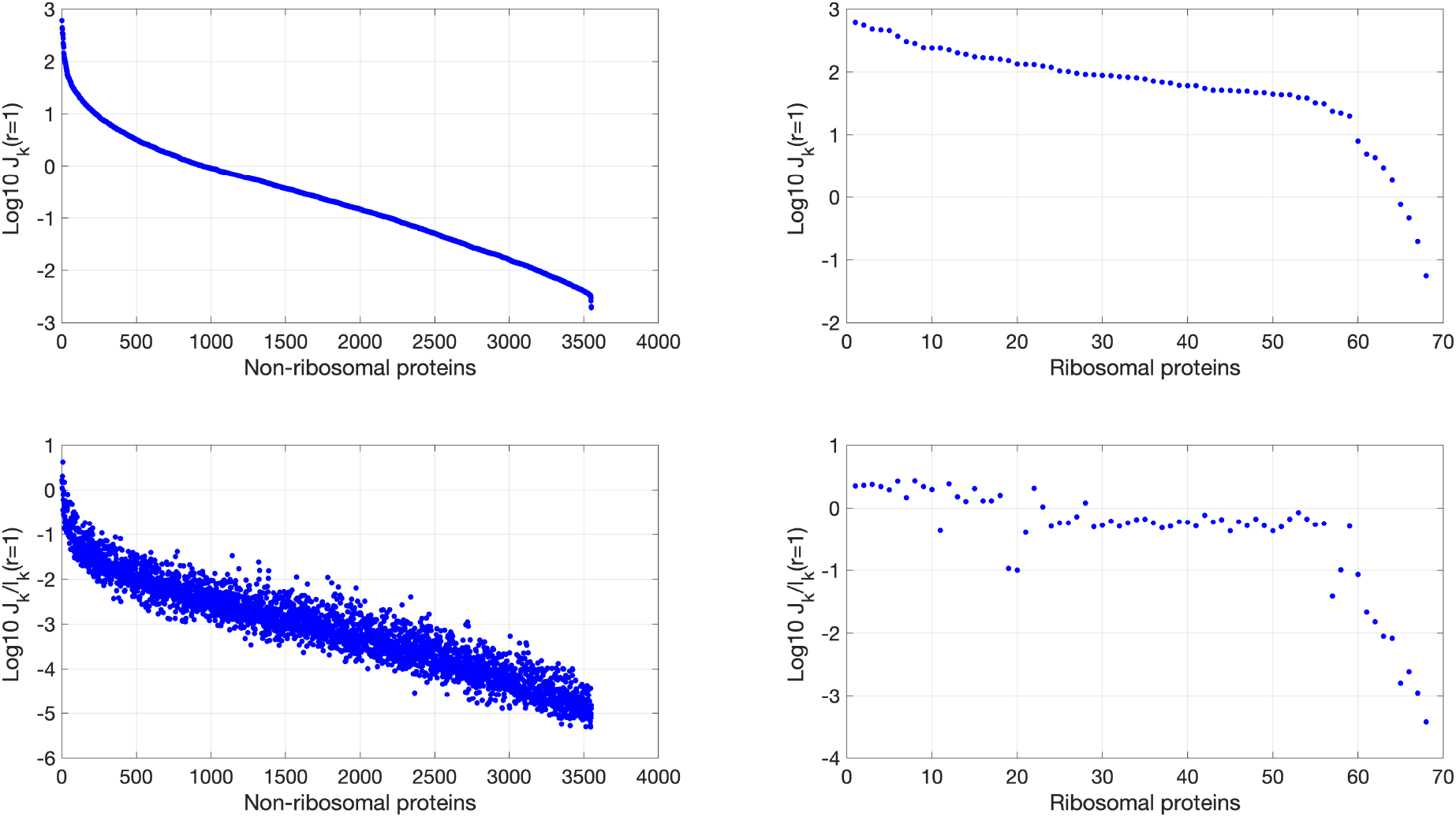
(Top:) Maximum resources recruitment strength *J_k_*(*r* = 1) for the set of non-ribosomal (left) and ribosomal protein expression genes (right) sorted by magnitude in logarithmic scale. (Bottom:) Ratio between the sorted maximum resources recruitment strength *J_k_*(*r* = 1) and the corresponding protein length (aa).

Not all genes are expressed all the time. As a proxy to estimate how many genes are active at a given time we calculated the cumulative sum of the maximum resources recruitment strength and obtained how many genes being expressed are required to explain both 95% and 99% of the total cumulative sum. We did this independently for both ribosomal and non-ribosomal proteins. Figure 7 shows the results obtained. *E coli* has around 4225 protein-coding genes [28, 22]. We had information obtained from [19] to get the values of *J_k_*(*r* = 1) for 3551 non-ribosomal and 68 ribosomal ones. That is 3619 genes, around 86% of all *E coli* genes. Therefore, the results obtained are enough representative of the cell.

**Figure 7:**
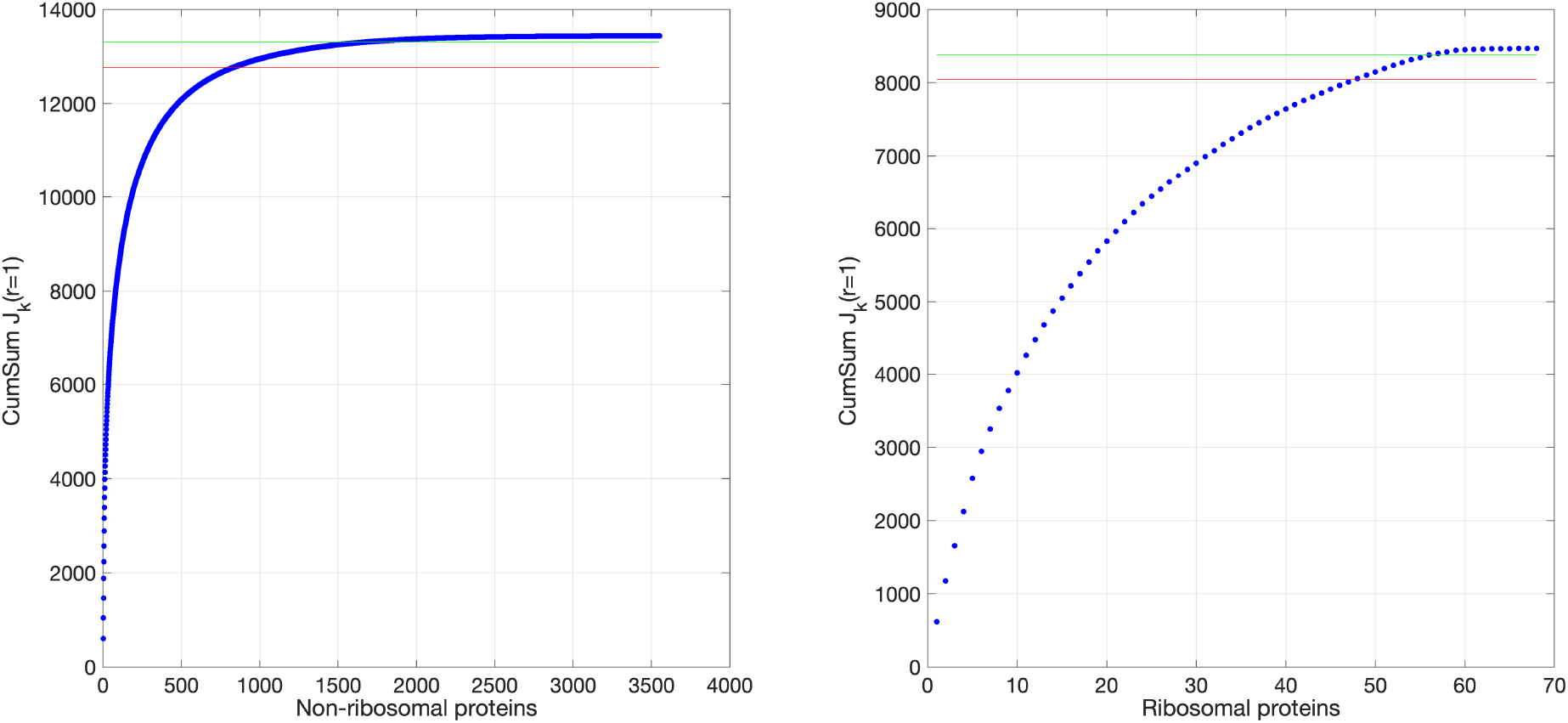
Cumulative maximum resources recruitment strength for the set of non-ribosomal (left) and ribosomal protein expression genes (right). The horizontal lines correspond to the 95% (red) and 99% (green) levels respectively.

Our results show that out of the 68 ribosomal genes, 49 of them (72%) explain 95% of the cumulative sum of the maximum resources recruitment strength of the ribosomal genes. To explain 99% we need 57 ribosomal genes (84% of them). On the other hand, for non-ribosomal genes we need 875 out of 3551 genes (25%) to explain 95% of the cumulative sum and 1735 (49%) to explain the 99%. These results show again that ribosomal genes are continuously needed for the cell and thus are continuously expressed. On the contrary, non-ribosomal genes are not required at all times. Since transcription and translation are energetically expensive processes, these genes are regulated to be expressed only when required. This explains the very low values obtained for the resources recruitment strength for most of them. This reflects the fact that most of the time they are “switched-off”.

The results above suggest that to get a reliable average value of the resources recruitment strength *J*_avg_ of a gene as a function of the number of free ribosomes *r* we must consider, on the one hand ribosomal and non-ribosomal genes as two differentiated sets and, on the other, we must consider some estimation of the number of active genes. To this end, a conservative estimation will consider as active the number of genes that explain 99% of the total cumulative sum of the maximum resources recruitment strength. Thus, we used the estimation of the sum 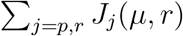 obtained in Section 13 (see Figure 5(left)) and assumed that the fraction of 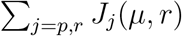 allocated for ribosomal and non-ribosomal proteins keep constant with the growth rate. Then, using the value of this fraction obtained from the data [19] for *t_d_* = 21.5 minutes, we considered 57 ribosomal and 1735 non-ribosomal active genes respectively to get the average resources recruitment strength *J*_avg_ for both ribosomal and non-ribosomal genes.

Figures 8 and 9 show the values obtained for *J*_avg_(*μ, r*) for an *E. coli* protein as a function of the number of free ribosomes *r* and both the quantity of available mature ribosomes *r_a_* and the duplication time *t_d_* respectively.

**Figure 8:**
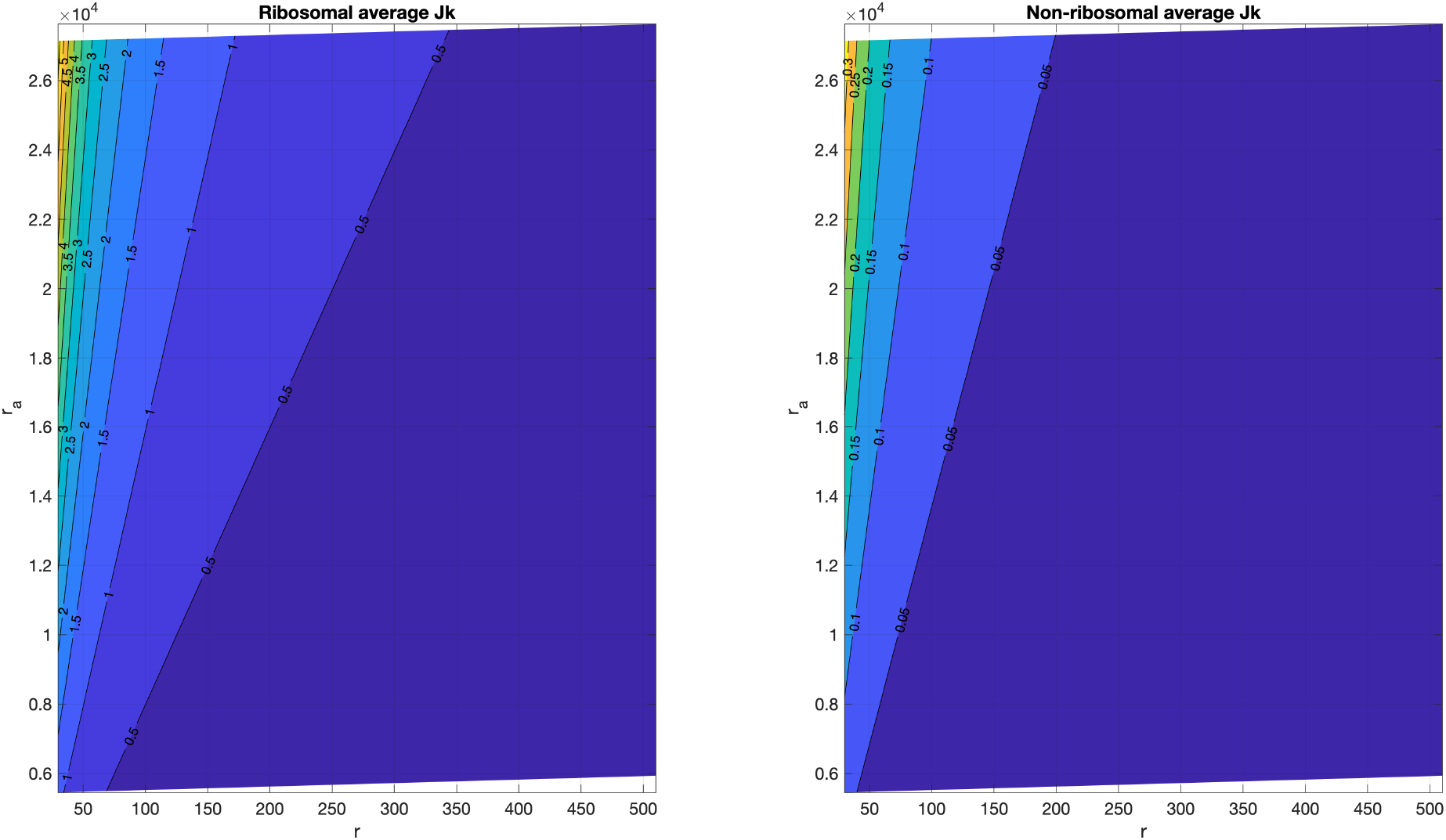
Estimated values obtained for *J*_avg_(*μ, r*) for an *E. coli* protein as a function of the number of free ribosomes *r* and that of available mature ribosomes *r_a_*. (Left:) ribosomal proteins. (Right:) non-ribosomal proteins.

**Figure 9:**
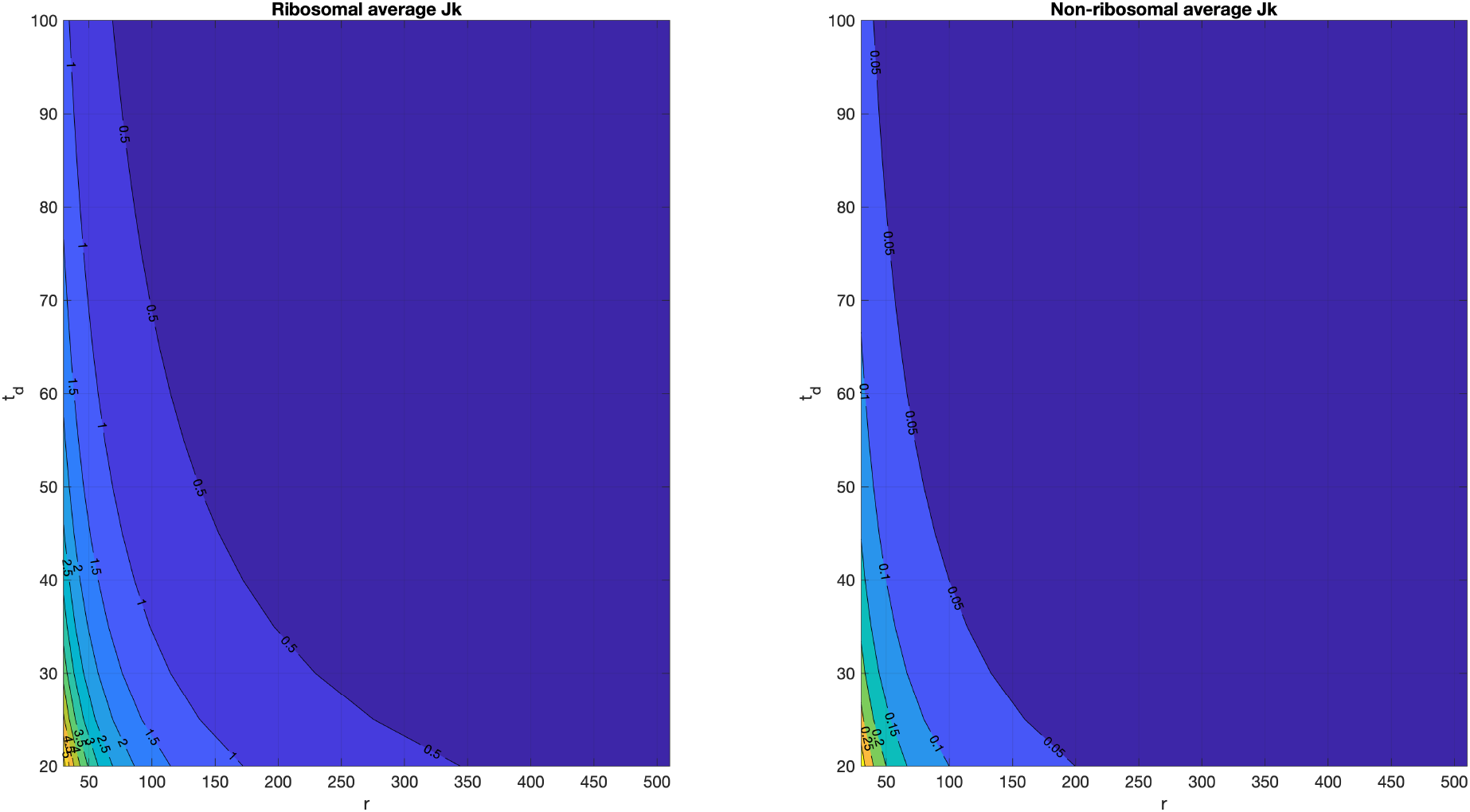
Estimated values obtained for *J*_avg_(*μ, r*) for an *E. coli* protein as a function of the amount of free ribosomes *r* and the duplication time *t_d_*t. (Left:) ribosomal proteins. (Right:) non-ribosomal proteins.

## 15. Estimation of the number of free ribosomes in the cell

Estimation of the number of free ribosomes in the cell, *r*, is key for assessing the competition among the cell circuits for cellular resources. The results in the previous sections suggest that an extremely low number of free ribosomes, with order of magnitude in the range 10^1^, gives a high sensitivity of the total cumulative sum of the maximum resources recruitment strength with respect to variations in the amount of free ribosomes (see Figures 4 and 5) while an order of magnitude in the range 10^2^ is enough to keep a robust value of the total amount of recruited resources with respect to variations in the number of free ribosomes. That is, by not expressing superfluous resources, the cell forces a competition for them that induces a high sensitivity of the total amount of recruited resources with respect to variations in the number of free ribosomes, while a small surplus of superfluous resources induces robustness.

To evaluate the range of expected values of *r* we used experimental data of the translation efficiency per mRNA. Notice from the dynamics (19) for a protein *p_k_* we can define:

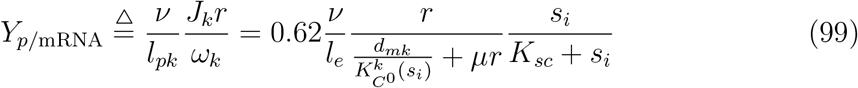

where *s_i_* is the copy number of molecules of intracellular substrate, *d_mk_* is the mRNA degradation rate, recall 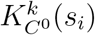 is a substrate dependent parameter essentially related to the RBS strength and we consider that substrate availability will only affect translation and not transcription (see Section 2). Notice *Y*_*p*/mRNA_ is the copy number of protein produced per amount of transcript.

We used the data from [19] to estimate an upper bound for the number of free ribosomes *r* using (99) and the values for *Y*_*p*/mRNA_ obtained from expression (94):

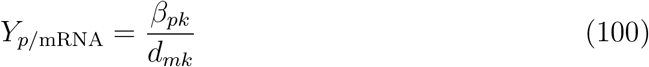

Then, the relationship between the RBS strength-related term 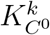 and the free ribosomes *r* becomes:

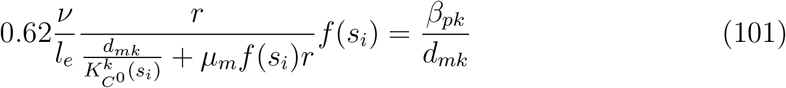

where *f*(*s_i_*) = *s_i_*/(*K_sc_* + *s_i_*) and we have used equations (55), (56) and (57) relating the specific growth rate *μ* with the maximum one *μ_m_* and the availability of intracellular substrate. Notice that, for any given protein and intracellular substrate availability, the number of free ribosomes will determine the required value of the RBS strength-related term 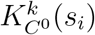 to attain the experimental value of the translation efficiency per mRNA *Y*_*p*/mRNA_.

The translation efficiency given by expression (101) depends on the ribosomes density 1/*l_e_*. An average ribosomes density around 4.2 ribosomes per 100 codons in optimal growth conditions has been reported in the literature for the prokaryote *L. lactis* [30]. This value is the same we obtained from the data in (94) by considering the total number of available active ribosomes Φ*_b_r_a_* obtained in Section 14 (see Figure 5) and dividing it by the sum of the lengths of all proteins weighted by a factor 0.5 to account for the estimation that 50% of the genes are active (explain 99% of the cumulative sum of the maximum resources recruitment strength). Similar values are found for other organisms [14]. For *E. coli* a value of 3.5 is given in [28]. The ribosomes density is inversely log-linearly related to the length of the coding sequence, with a slope quite consistent for a variety of organisms [14]. To account for this, we approximated a power law consistent with the findings in [14] and resulting in 4 ribosomes per 100 codons for an average protein length of 330 codons. We obtained the relationship:

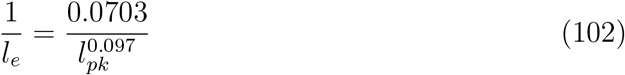

This gives a range *l_e_* ⊂ [18, 31], with a value *l_e_* = 25 for the average protein length. The minimum value is consistent with the shortest protein length (18 codons) in the database we used.

Figure 10 shows the results obtained for the set of all ribosomal and non-ribosomal proteins and their average values.

**Figure 10:**
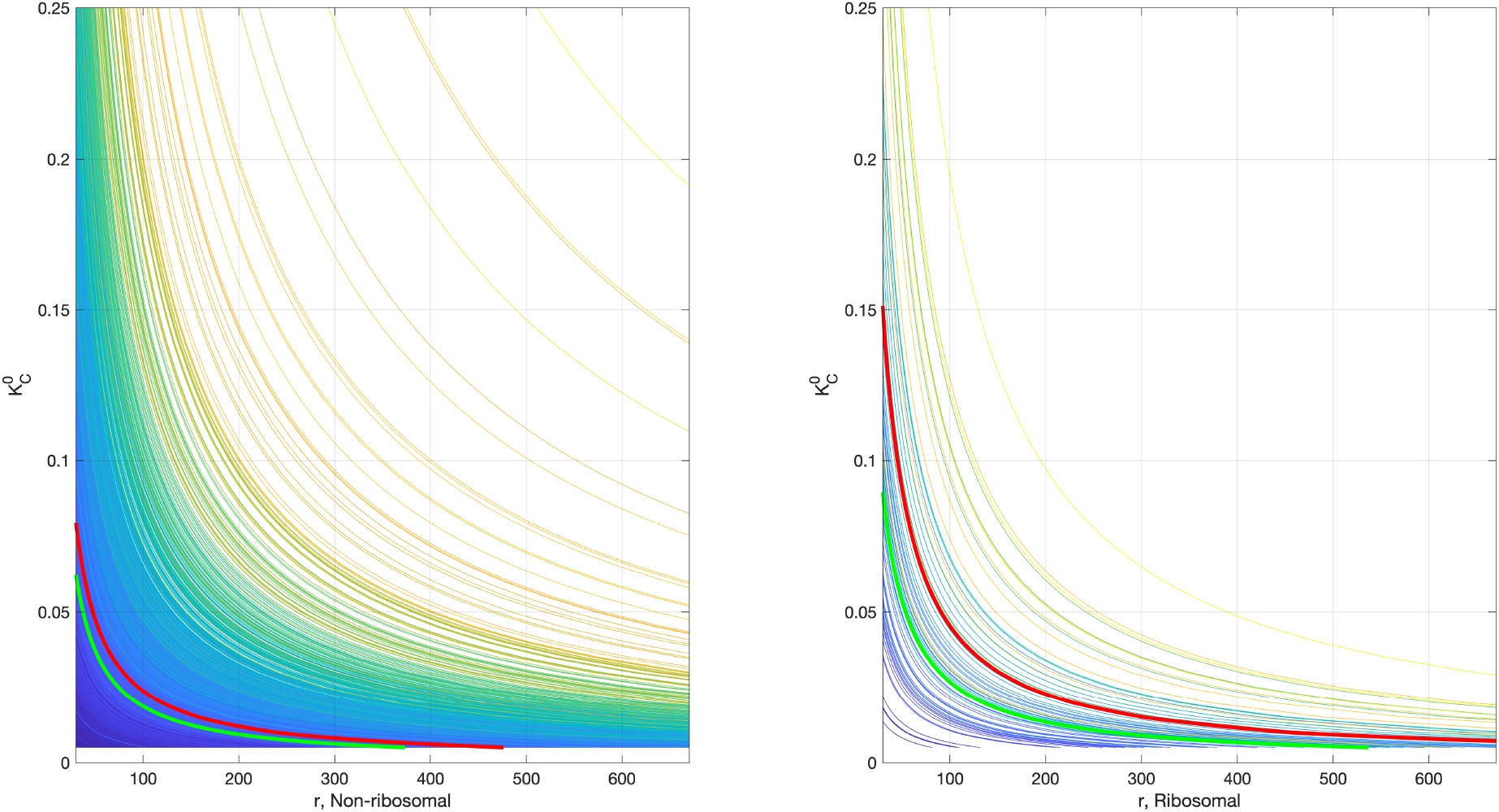
Relationship between the RBS strength-related term 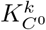 and the number of free ribosomes *r* obtained using the experimental data in [19] for non-ribosomal (left) and ribosomal (right) proteins in *E. coli*. Thin lines correspond to the experimental value of the translation efficiency per mRNA *Y*_*p*/mRNA_ for each protein. The red thick line corresponds to the mean for all proteins in the corresponding non-ribosomal and ribosomal sets. The green thick line corresponds to the approximated mean when the term associated to the maximum specific cell growth rate is neglected in the expression (101).

From the results shown in Figure 10, notice that the number of free ribosomes *r* required to explain the experimental value of the translation efficiency per mRNA *Y*_*p*/mRNA_ for each protein increases as the value of the RBS strength-related term 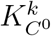 decreases. Indeed, the more free ribosomes are available, the less competition for shared resources. The number of free ribosomes *r* is an indicator of the level of competition for resources. Thus, expression (101) implies that a gene producing short-living transcripts will require, for the same level of competition, a stronger RBS to achieve the same translation efficiency per mRNA *Y*_*p*/mRNA_ as one with long-living transcripts.

To estimate an upper limit for the copy number of free ribosomes required to achieve the experimental translation rates per mRNA, we considered an upper bound for the RBS strength-related term 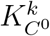.

Recall from equation (10) that 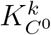 term is a function of the intracellular substrate availability. Since the data we used from [19] was obtained for fast growing cells, we can consider intracellular substrate saturation. Under this condition we get:

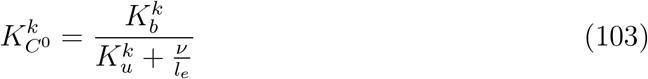

Using the value of *ν* in Table 12 and the range of *l_e_* given above, we estimate 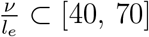 (molec^−1^ · min^−1^). On the other hand, the values of the association and dissociation rates of the ribosome to the RBS, 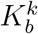 and 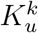, may vary in a large range. Values 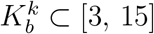 (molec^−1^ · min^−1^) are found in the literature (see Table 12). We use a conservative upper bound 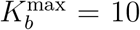 (molec^−1^) considering binding is diffusion controlled. From the literature, we consider a range for the dissociation rate 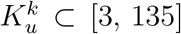 (min^−1^). Overall, these estimates give us a range (under the assumption of intracellular substrate saturation) 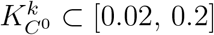 (molec^−1^).

From the results shown in Figure 10, notice that a maximum number of free ribosomes *r* ≈ 350 can confidently explain the translation efficiencies per mRNA *Y*_*p*/mRNA_ for almost all proteins while maintaining the value of the RBS strength-related term 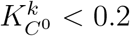. This estimation for the amount of free ribosomes is in complete agreement with the estimation *r* = **N**(350, 35) in [23].

With the upper limit *r* ≈ 350 we could explain the translation efficiencies per mRNA *Y*_*p*/mRNA_ calculated as 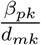 (see equation (101)) using the data in [19] but for a small set of 80 non-ribosomal proteins out of 3551 (2.25%). In 52 of them, this could be attributed to their extremely long-living transcripts. In the remaining 28 ones, to their very high translation efficiency per mRNA *Y*_*p*/mRNA_ expected from their values of *β_pk_* and *d_mk_* given in [19]. This can be explained by rewriting (101) as:

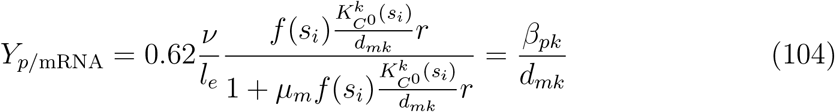

Figure 11 shows a plot of the function (104), as a function of its argument 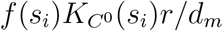 for two mRNA degradation rates corresponding to short and long-living mRNAs and ribosomes densities in the range *l_e_* = [18, 31]. Notice 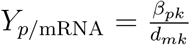 saturates at the maximum attainable value:

**Figure 11:**
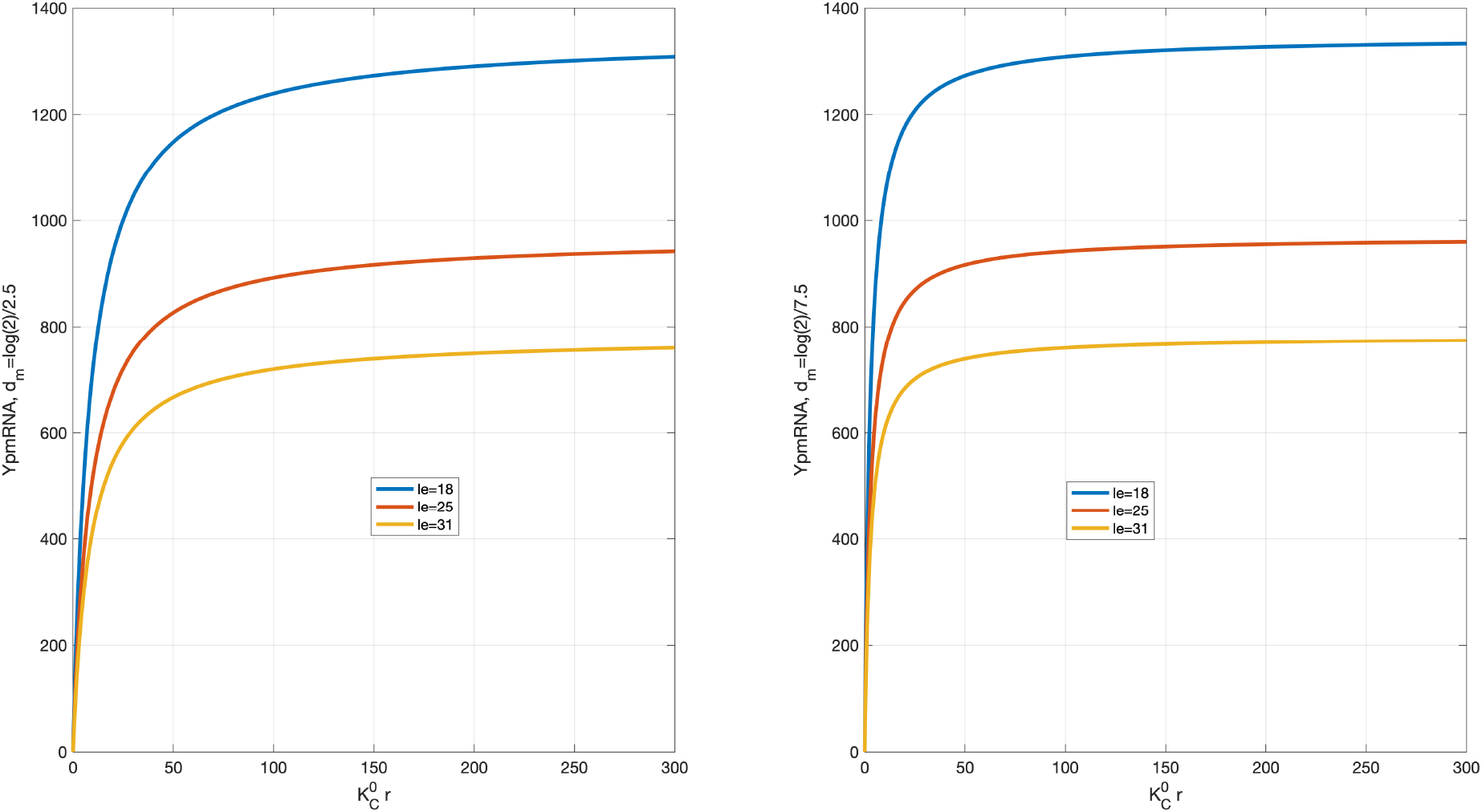
Translation efficiency per mRNA *Y*_*p*/mRNA_ as a function of *f*(*s_i_*)*K*_*C*^0^_*r* for two mRNA degradation rates corresponding to mRNA half-lives 2.5 minutes (left) and 7.5 minutes (right) for ribosomes densities corresponding to *l_e_* = {18, 25, 31} codons.

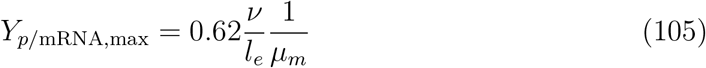

Saturation is reached for lower values of the argument 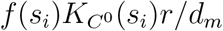 as the transcripts have longer half-lives, i.e. smaller values of the degradation rate *d_m_*.

The few cases our model could not predict all have values of 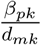 above *Y*_*p*//mRNA,max_ for the values of *v* and *l_e_* used in the model.

On the other hand, it is interesting to notice that for very low values of *f*(*s_i_*)*K*_*C*^0^_(*s_i_*)*r*/*d_m_* such that 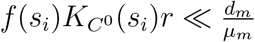 we can approximate:

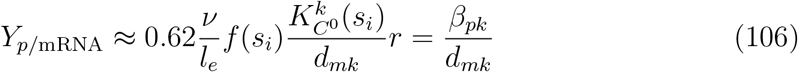

from which we get:

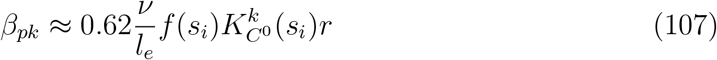

That is, for a highly competitive scenario where the number of free ribosomes is sufficiently small (e.g. in the order of few tens to few hundreds for typical values of *d_mk_*, *μ_m_* = 0.032 min^−1^ and *f*(*s_i_*)*K*_*C*^0^_(*s_i_*) at its maximum estimated value *f*(*s_i_*)*K*_*C*^0^_(*s_i_*) = 0.2) the translation rate (proteins per mRNA per time unit) is proportional to the ribosomes density 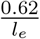, the effective maximum translation rate per codon attainable for a given substrate availability *vf*(*s_i_*), the RBS strength 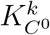, and the available free ribosomes *r*.

Notice under this scenario the translation rate will suffer large stochastic fluctuations caused by stochastic fluctuations in the number of free ribosomes. In this case, the transcription rate for a given RBS-strength mainly depends on the competition for cellular resources and, therefore, on the number of free ribosomes *r*, and it is largely independent of the specific growth rate.

## 16. Sensitivity of the resources recruitment strength to RBS and promoter can explain the variation of the cell mass distribution with growth rate

The relative distribution of ribosomal and non-ribosomal protein mass in the cell depends on the cell growth rate, so that the ribosome content increases linearly with growth rate [5, 35, 27, 9]. Existing resource allocation models explain this as a result of optimal allocation of cell resources between the ribosomal and protein (non-ribosomal) fractions balancing the demands of protein synthesis and nutrient influx under the constraint that the sum of both fractions keeps constant [35].

We used the data in [5] of the ribosomal and non-ribosomal protein mass fractions as a function of growth rate and the corresponding effective translation rates to check whether our model is able to describe the linear increase of ribosomes content with growth rate.

In our model, the relative resources recruitment strength of a given protein equals its relative mass fraction in the cell at steady state, according to equation (72). Therefore, the relative distribution of ribosomal and non-ribosomal protein mass must be reflected in their relative distribution of resources recruitment strengths.

As a first step, we considered the lumped resources recruitment strengths (73)–(74) and estimated the RBS-strength related parameters 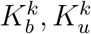, the transcription rates *ω*_k_ with *k* = {*r*, wt} and the fraction Φ_*t*_ so that our model provides a good fit of the specific growth rate at steady state. Notice that we did not estimate the parameters in *J_r_*(*μ, r*), *J*_wt_(*μ, r*) to try and directly fit the experimental relative distribution of resources recruitment strengths. Instead, we were interested in checking whether a good fit of the specific growth rate implies ribosomal and non-ribosomal resources recruitment strengths such that their relative values fit the experimental ones. This, in turn, implies fitting the relative mass fractions in the cell.

We used the model expressions at steady state in Section 10 and the only input information given to the model was the values of the effective maximum translation rate *v_t_* as a function of growth rate obtained from [5]. We ran 200 instances of the parameter optimization using the optimization global optimization software MEIGO [12] (available at http://gingproc.iim.csic.es/meigo.html) and obtained the weighted mean of the 25 runs achieving the best minimum value for the sum over the experimental data points of the absolute growth rate prediction error. We considered *N_r_* = 57; and *N*_wt_ = 1735, corresponding to the number of genes that explain 99% of the cumulative sum of the resources recruitments strengths for ribosomal and non-ribosomal proteins respectively (see Section 15). We also considered the average mRNA degradation rates *d_m,r_* = 0.16 (1/min) and *d*_*m*,wt_ = 0.2. The resulting average best fit estimated parameters are given in Table 3.

**Table 1:** Phenomenological models relating cell and protein dry mass and specific growth rate and their best fit parameters for the data in [5].

| Model | Model parameters |  |  |
| --- | --- | --- | --- |
| $m_c = \bar{m}_0 e^{\bar{\gamma}\mu}$ | $\bar{m}_0 = 123.022 \cdot 10^{-15} \text{ (g)}$ | $\bar{\gamma} = 68.5547 \text{ (min)}$ | |
| $m_p = \bar{m}_{p0} e^{\beta\mu}$ | $\bar{m}_{p0} = 77.3748 \cdot 10^{-15} \text{ (g)}$ | $\beta = 61.7813 \text{ (min)}$ | |

**Table 2:**
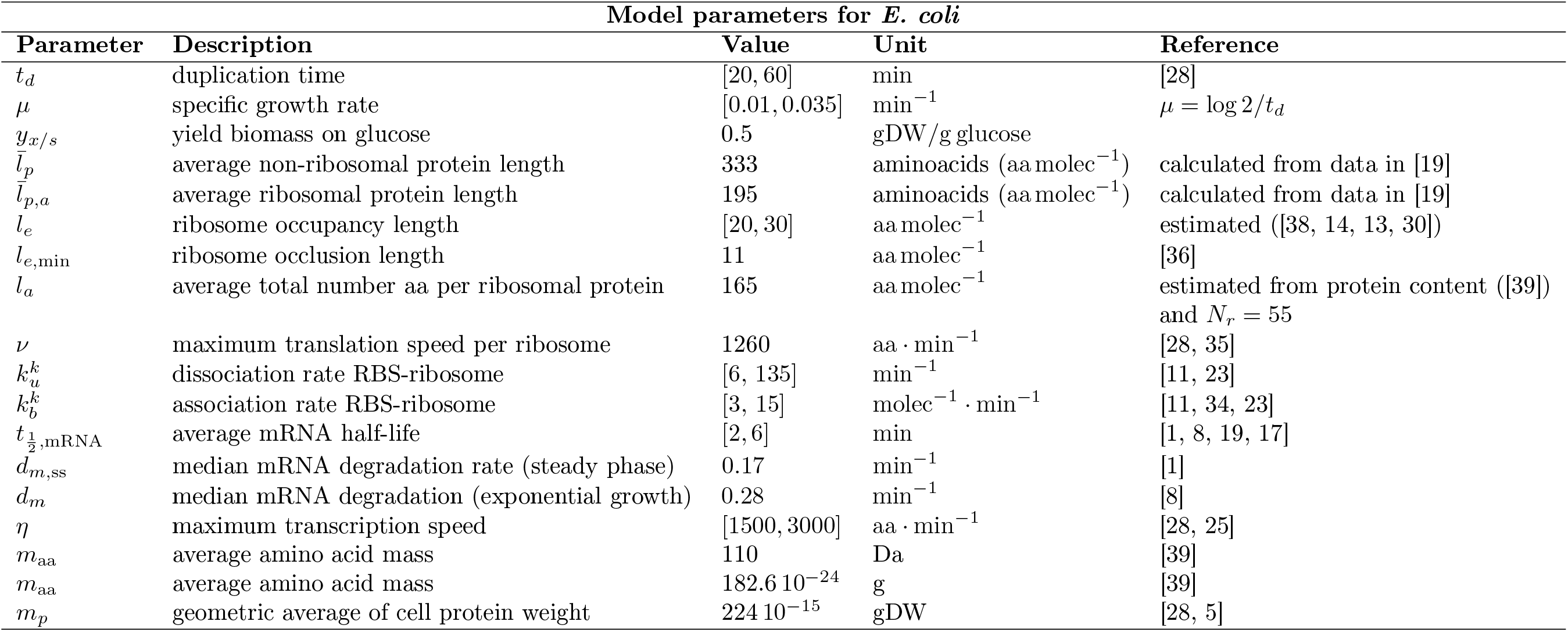
Model parameters.

**Table 3:** Average best fit estimated parameters of the RBS-strength related parameters 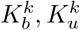, the transcription rates *ω*_k_ with *k* = {*r*, wt} and the fraction Φ_*t*_ of mature ribosomes with respect to the total number of ribosomes.

| $K_u^r$ | $K_u^{nr}$ | $K_b^r$ | $K_b^{nr}$ | $\omega_r$ | $\omega_{nr}$ | $\Phi_t$ |
| --- | --- | --- | --- | --- | --- | --- |
| 130.74 | 3.11 | 5.81 | 11.80 | 5.36 | 0.028 | 0.78 |

Figure 12 shows the results of the optimization. Notice the good agreement between the experimental and the estimated growth rate. Notice also that the estimation for the maximum number of free ribosomes *r* = 350 used in previous sections is consistent with model results for cells growing up to some *t_d_* = 25 minutes (*μ* ≈ 0.028. For faster growing cells, the number of free ribosomes much increases. Notice though, that also the total number of ribosomes (both experimental and estimated) much increases for very fast growing cells. Thus, the fraction of free ribosomes with respect to the total number only increases from 0.08% up to 1.37% for cell duplication times between 100 and 24 minutes respectively even though the number of free ribosomes multiplies by almost 200-fold (see Figure 13). Notice also the logarithmic affine relationship between the number of free ribosomes *r* and its flux *μr*, (log_10_(*r*) ≈ 4.11 + 0.79log_10_(*μr*)) reflecting a power-law relationship between growth rate and number of free ribosomes.

**Figure 12:**
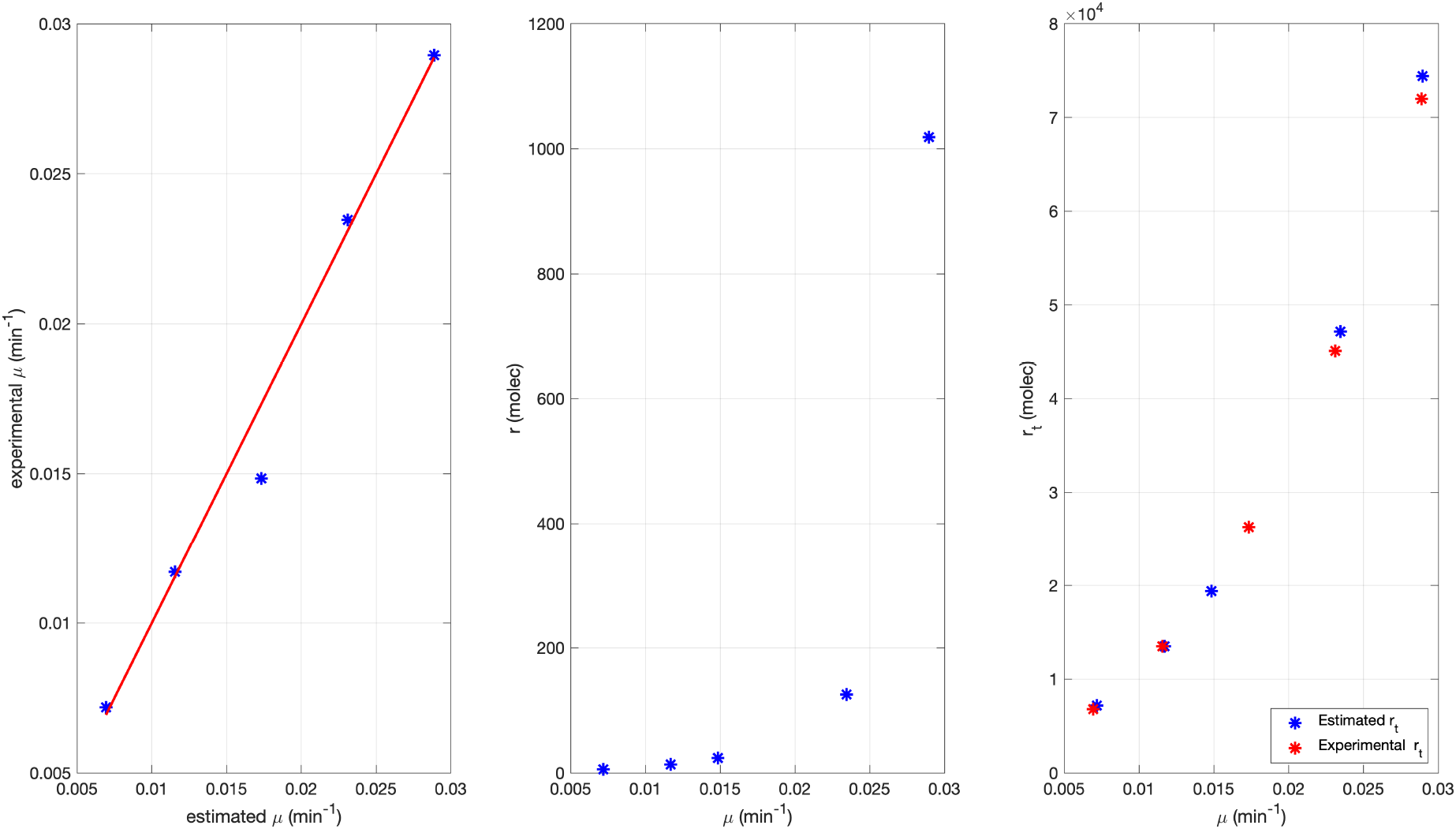
Estimated versus experimental growth rate (Left). Estimated number of free ribosomes (Center). Estimated and experimental number of total ribosomes (Right)

**Figure 13:**
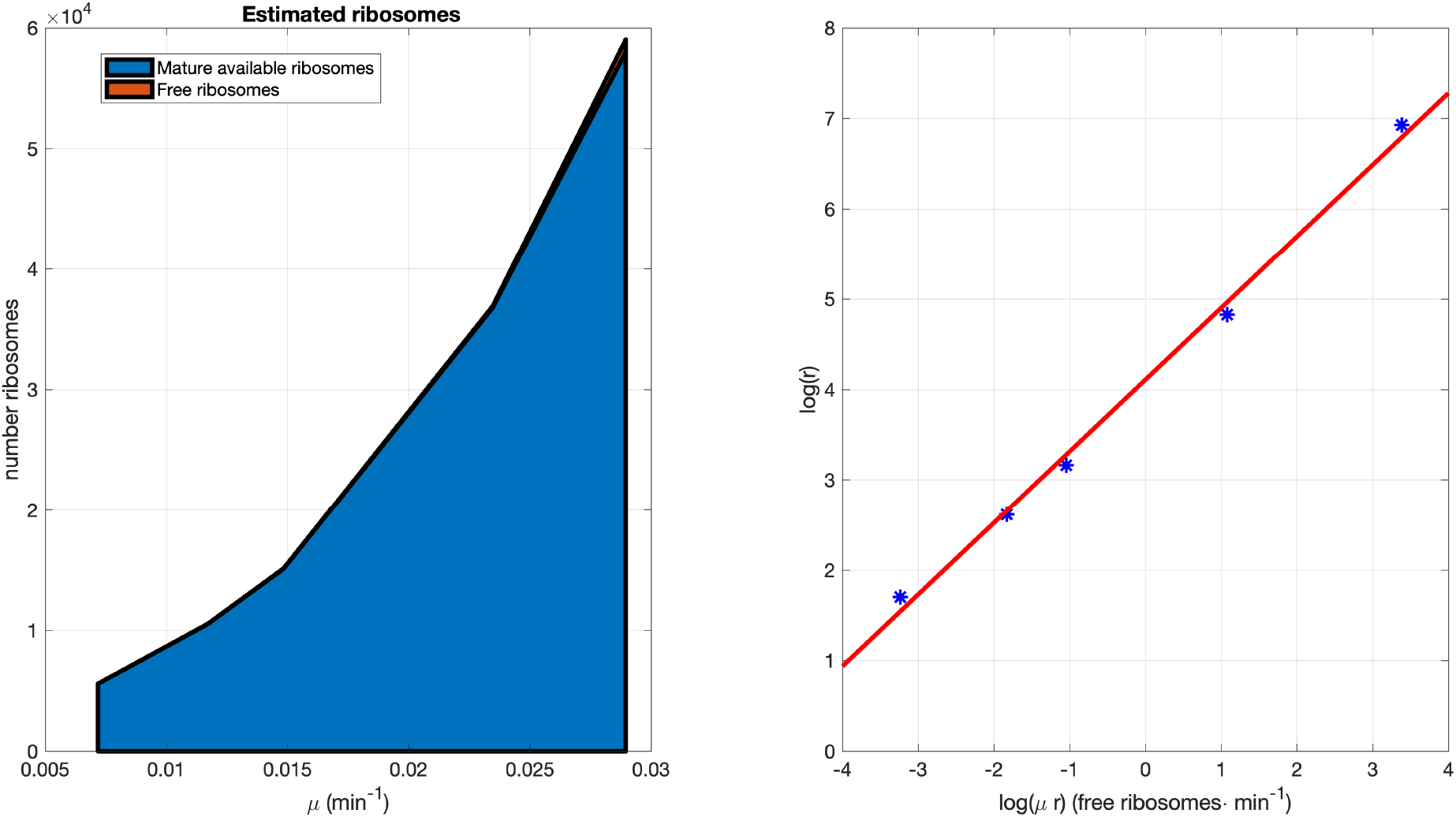
Estimated number of free and total ribosomes (Left) and logarithmic affine relationship between the number of free ribosomes *r* and its flux *μr* (Right)

Notice the estimated value of the fraction Φ_*t*_ of mature ribosomes with respect to the total number of ribosomes. is very close to the average experimental value Φ*_t_* = 0.8 [4, 5]. Notice also the clear difference between the parameters corresponding to ribosomal proteins as compared to those of non-ribosomal ones. The results suggest that the ribosomal RBSs are much weaker than the non-ribosomal ones. Thus ribosomes have much less affinity for the ribosomal RBSs than for non-ribosomal RBSs. Figure 14 shows the estimated translation initiation rate *k_e_* and the estimated effective RBS strengths 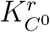 and 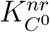 as a function of the specific growth rate *μ*. As seen, the effective RBS strengths of non-ribosomal proteins is much higher than that of the ribosomal ones. Notice though, that as the growth rate increases −tantamount in our model to increasing intracellular substrate *s_i_*− the ribosomal effective RBS strength 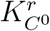 keeps almost constant (with a slight decrease around 12%) while the non-ribosomal one decreases by almost a 40%. Notice also the clear difference between the terms 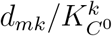 for both ribosomal and non-ribosomal proteins when as a function of the flux of free resources *μr* (cf. the expression of the resources recruitment strengths given in equations (73)–(74)). The ribosomal proteins keep much higher values of 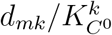 for all values of the flux of free resources *μr*. This, considering the monotonous increasing power-law relationship between the growth rate and the number of free ribsosomes, will imply that the value of the ribosomal resources recruitment strength *J_r_*(*μ, r*) will decrease much slower than that of the non-ribosomal proteins as the growth rate increases. The plots in Figure 15 confirm this.

**Figure 14:**
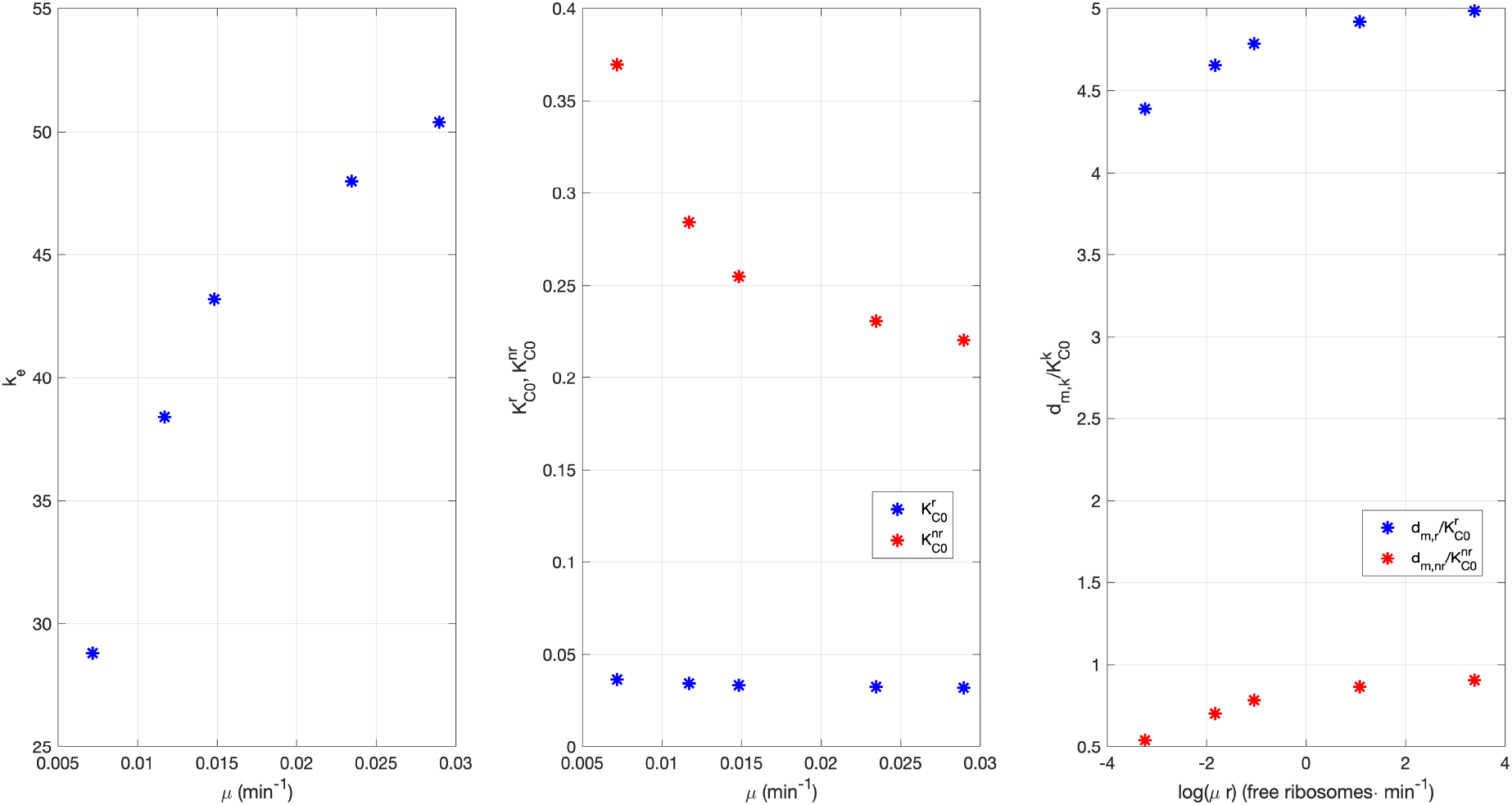
Estimated translation initiation rate *k_e_* (Left) and estimated effective RBS strengths 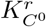 and 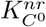 (Center) as a function of the specific growth rate *μ*. (Right): plot of 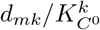 as a function of the flux of free resources *μr*. A log scale has been used.

**Figure 15:**
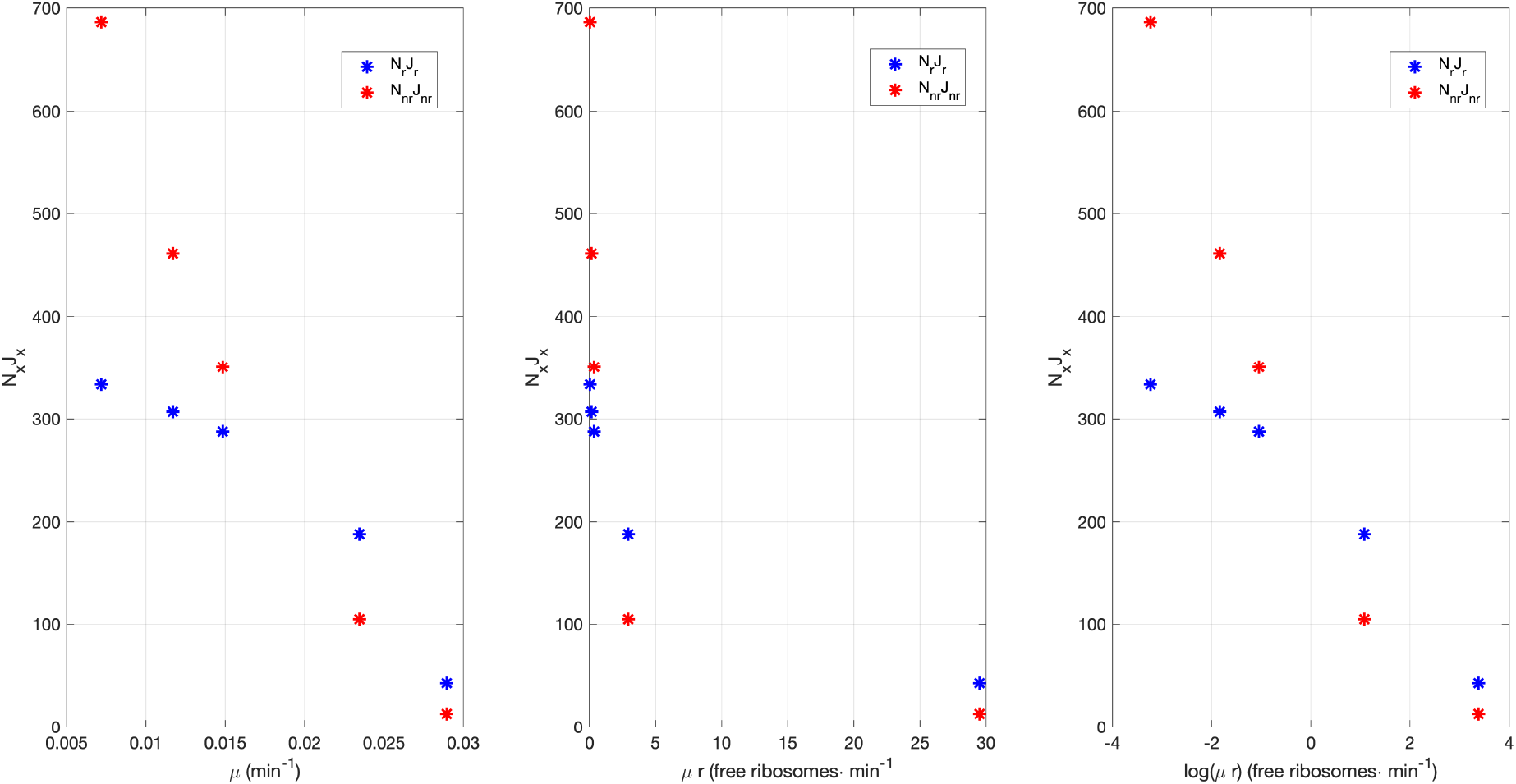
Estimated total resources recruitment strengths *N_r_ J_r_* and *N_nr_ J_nr_* as a function of growth rate *μ*(Left) and the free resources flux *μr* using linear (Center) and logarithmic scales (Right).

The differential behaviour between the ribosomal and non-ribosomal resources recruitment strengths is behind the differential mass distribution between ribosomal and non-ribosomal cell protein content as growth rate increases. Thus, the evaluation of the mass fractions using (79)–(80) provides the results shown in Figure 16 showing a very good agreement between the experimental values and the ones provided by the model.

**Figure 16:**
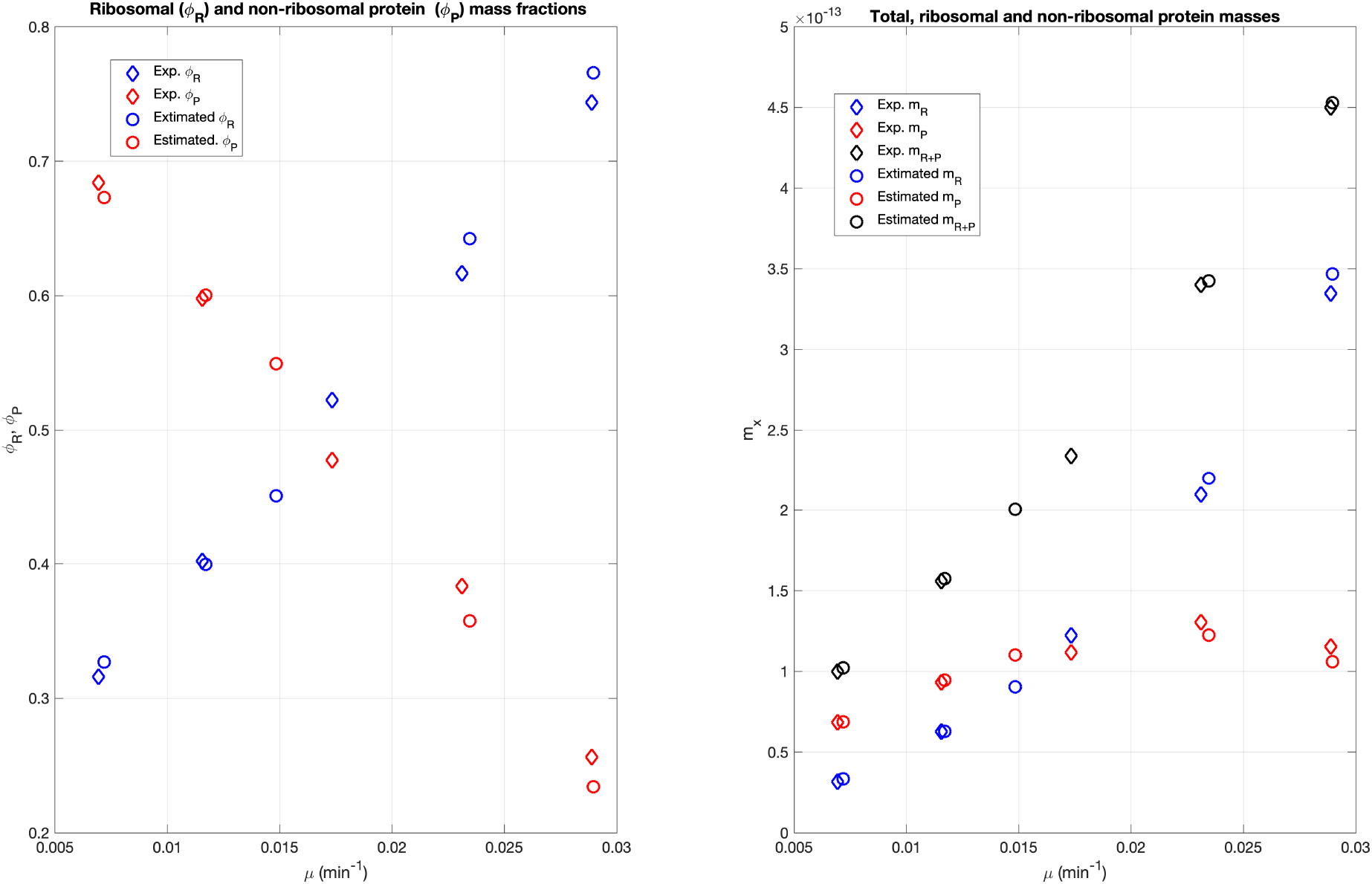
Estimated versus experimental mass fractions of ribosomal and non-ribosomal proteins in *E. coli* (left) and their absolute values along with the total protein cell mass (right).

Notice that the results above suggest that to have a robust expression rate as the growth rate increases, the optimal strategy consists of having a weak RBS and a strong promoter. Interestingly, the values we obtained for the transcription rates are in the same order of magnitude as the mean values obtained from the data in [19] −*ω_r_* = 2.4 and *ω_nr_* = 0.05 respectively− and show a much higher value for the average lumped ribosomal protein we considered than for the non-ribosomal one. The optimal productivity will depend on the trade-off between RBS and promoter strengths and their effect on growth rate, as analysed in the next sections.

## 17. The differential role of RBS and promoter strengths

It is well known that varying combinations of transcription and translation rates affect the stability of metabolic networks [29] and the trade-off between desired expression levels and noise [19] and between expression of endogenous and synthetic genes and growth [43, 16].

We were interested in analysing the differential role of RBS and promoter strengths for endogenous proteins, that is, for the host ones, assuming there is no loading effect.

To this end, we analysed the protein abundance of an endogenous protein as a function of the of the expression space by varying the transcription rate *w_k_*, and the RBS affinity for the mRNA (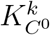 in our model). We are now focused on understanding the role of promoter and RBS strengths. Loading effects (that would be expected if overexpressing an exogenous protein, as considered in Section 18) will introduce interactions that will hide the differential effects of varying the transcription rate and the RBS affinity for the mRNA. To avoid this, we considered the case of expression of a protein *A* under the assumptions that 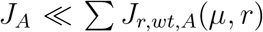 and 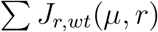 is constant (c.f. equations (88) and (89) in Section 11). This assumption holds for very low values of *w_A_*. Under these conditions, the flux of free resources *μr* can be assumed to depend only on the constant value of *J_r,wt_*(*μ, r*).

It is important to note that *J_A_*(*μ, r*) does not directly define the expression of the protein *A*, that is, the steady state value of the protein mass 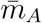. Recall from Section 9 (see also equation(88)) that it is the relative value of the resources recruitment strength *J_A_*(*μ, r*) with respect to the total sum *J_r,wt,A_*(*μ, r*) which defines the fraction of the cell protein mass *m_p_* that will be invested as protein *A*. Our assumptions imply constant 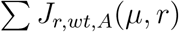 and constant flux of free resources *μr*. Under these conditions, from equation (85), the cell protein mass *m_p_* will also keep constant as a function only of the intracellular substrate *s_i_*. If in addition, we assume that the intracellular substrate *s_i_* keeps constant, then a variation in the value of *J_A_*(*μ, r*) will be monotonously related to a corresponding variation of 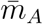.

Under the assumptions above, we can express:

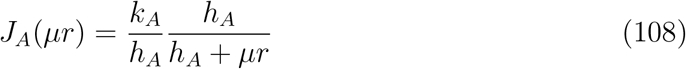

with:

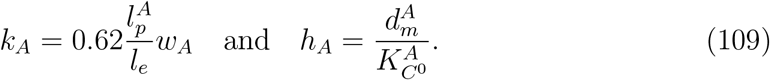

where *k_A_/h_A_* is the maximum value that *J_A_*(*μr*) can attain and *h_A_*/(*h_A_* + *μr*) is a Hill-like function which depends on the flow of free resources *μr* with *h_A_* as half-activation threshold.

Next we analyse the effect of increasing the transcription rate *w_A_* (for our purposes tantamount to increasing the transcription rate times the gene copy number) on the value of *J_A_*(*μ, r*). Recall we introduce changes in *w_A_* small enough so that our assumptions hold. As we vary *w_A_*, the term *k_A_/h_A_* will vary in the same proportion and *h_A_* will not be affected. Therefore, as the flow of free resources keeps constant under our conditions, the resources recruitment strength *J_A_*(*μ, r*) will vary in proportional relationship to the variations in *w_A_*. This, in turn, will affect in proportional relationship to the value of the mass 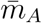. Therefore, under the condition of negligible metabolic load, variations in the transcription rate and the resulting variations in the amount of the expressed protein are proportional. Figure 17 (left) shows the agreement between this reasoning and the simulation results obtained with the full model.

**Figure 17:**
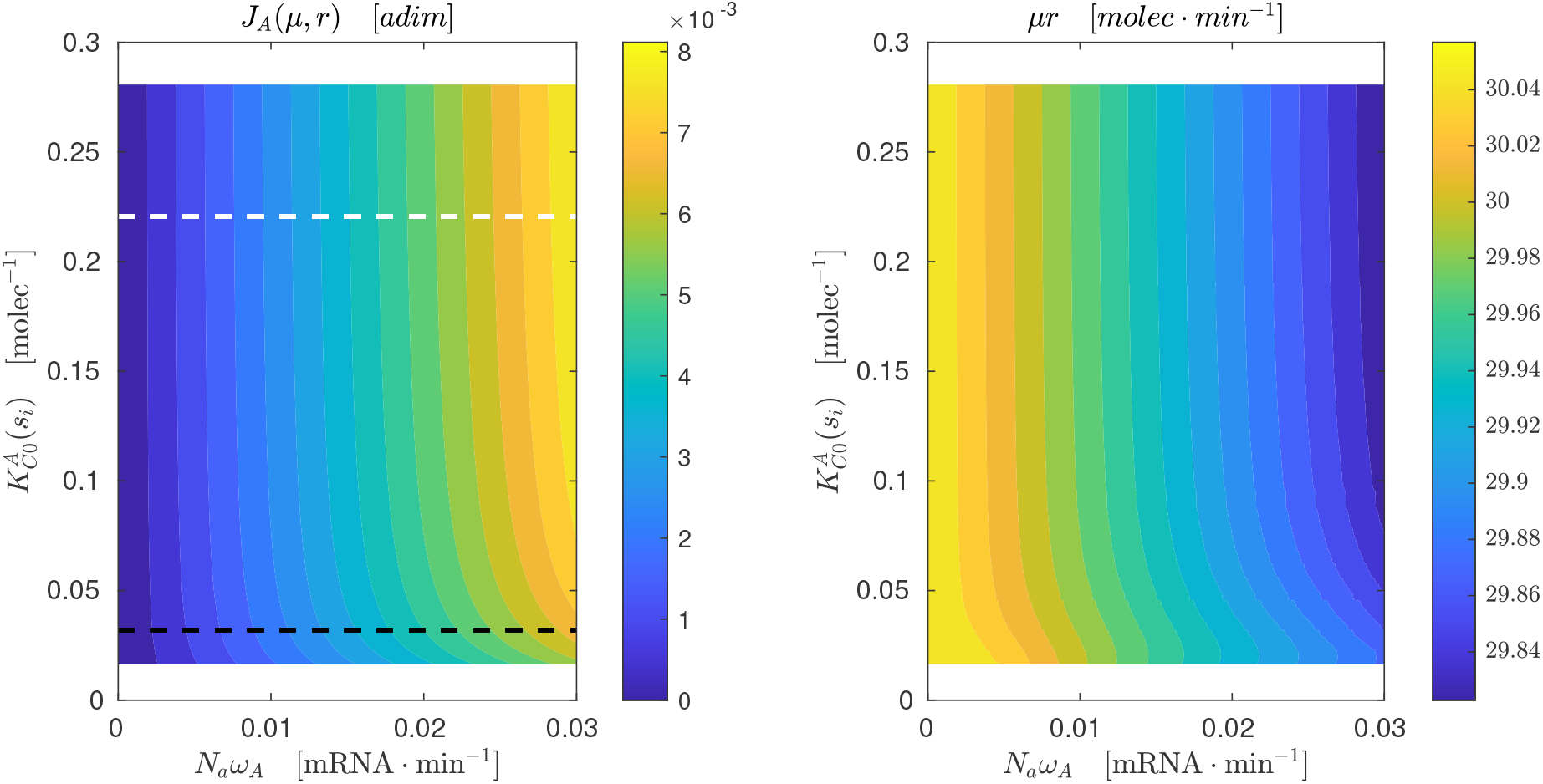
Effect of increasing transcription rate and RBS strength on the value of *J_A_*(*μ, r*) (left) and on the value of *μr* (right). The y-axis shows the full experimental range of 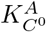 for *f*(*s_i_*) = 1, that is, saturated substrate. The average RBS strength value for *E. coli* ribosomal and non-ribosomal proteins are shown as black dashed and white dashed lines respectively. The x-axis shows the transcription rate values around the experimental average value of *E. coli* non-ribosomal genes (0.028min^−1^).

As for variations in the RBS strength, consider for instance the effect of increasing 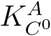, that is, increasing the RBS-ribosome association rate 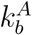 and decreasing the dissociation one 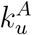 respectively. As we increase 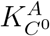, the term *k_A_/h_A_* will increase in the same proportion. Yet, the term *h_A_*/(*h_A_* + *μr*) will decrease. Which one of both opposite effects will determine the evolution of *J_A_*(*μr*) will depend on the relative weights of *h_A_* and the flux of free resources *μr*. If *h_A_* ≫ *μr*, an increment in 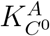 (compatible with the condition *h_A_* ≫ *μr*) will increase the steady state value of the protein mass *m_A_*, since *k_A_/h_A_* will grow and *h_A_*/(*h_A_* + *μr*) will remain mostly constant. However, if we keep 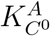 increasing we will reach a situation where *h_A_* ≪ *μr*. In the limit, equation 108 can be approximated as:

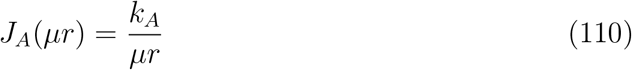

showing that at high RBS strength the resources recruitment strength *J_A_*(*μr*) saturates. When this saturation occurs, the protein mass 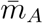 becomes independent of 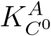. In summary, increasing the RBS strength has a limited effect to increase the expression of the protein *A*, saturating at high RBS strengths. Figure 17 confirms this general rule in the simulation results obtained with the full model.

In summary, RBS strength (together with mRNA degradation 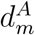) determines the sensitivity of the resources recruitment strength *J_A_*(*μ, r*) of a given protein *A* with respect to the flux of free resources *μr*. Thus, if protein *A* is expressed using a weak RBS, the value of *J_A_*(*μ, r*) will remain mostly constant with respect to variations of *μr*. On the contrary, if *A* is expressed with a strong RBS, the resources recruitment strength *J_A_*(*μ, r*) will decrease as *μr* increases. This result is confirmed in Figure 18 which shows the effect of varying *μr* on the value of *J_A_*(*μ, r*) as a function of the RBS strength.

**Figure 18:**
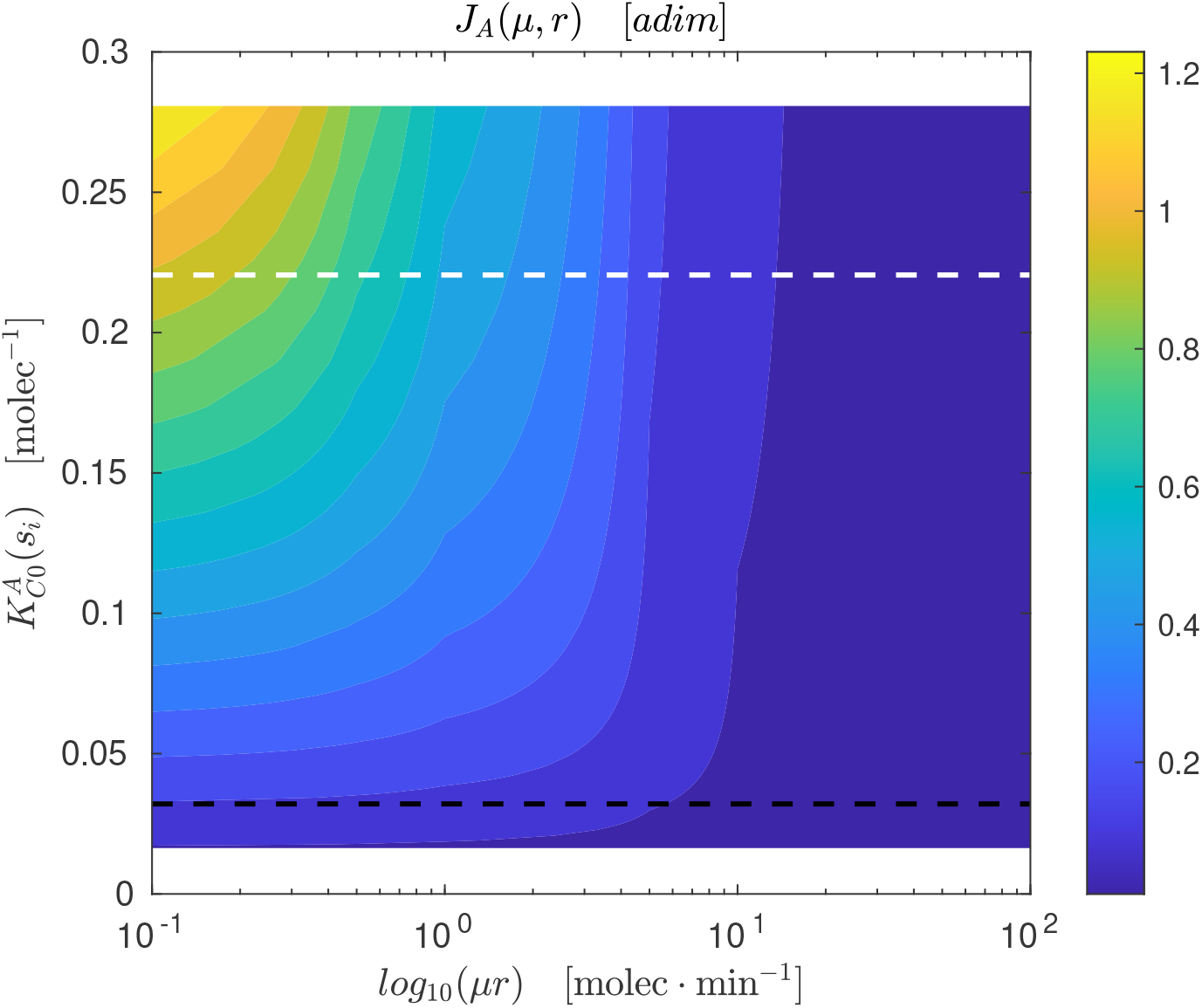
Effect of varying flux of free resources *μr* on the value of *J*(*μ, r*) with *k_A_* = 1 and 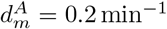. The y-axis shows the full experimental range of 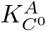 for *f*(*s_i_*) = 1, that is, saturated substrate. The RBS strenght value of ribosomal (black dashed) and non-ribosomal (white dashed) is shown. The x-axis shows a sweep in *μr* values for the range obtained in previous simulations.

As observed in Figure 18, there is a trade-off between increasing the expression of a protein by increasing the strength of RBS and increasing robustness with respect to *μr* by decreasing the RBS strength. It is not possible to achieve high expression and high robustness of the resources recruitment strength simply by adjusting the RBS strength.

This trade-off is consistent with the optimization results obtained in Section 16. Recall the ribosomal and non-ribosomal RBS parameters in Table 3, where ribosomal genes have low RBS strength (i.e. low 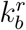 and high 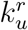) and high transcription rate, while the non-ribosomal genes have high RBS strength (i.e. high 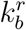 and low 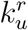) and low transcription rate. On the one hand, ribosomal genes have low RBS strength and high transcription rate, so their *J_r_*(*μ, r*) will remain mostly constant with respect to changes in the flow of free resources *μr*. On the other hand, non-ribosomal genes have high RBS strength and a low transcription rate. Therefore their expression will depend on the flow of free resources *μr*. As *μr* increases, the value of *J_nr_*(*μ, r*) will decrease. This difference in the values of RBS strength and transcription rate, long with the equivalence between the relative value of mass fractions *m_r,nr_* and the relative value of *J_r,nr_*(*μ, r*), allow us to explain the relative mass fractions distributions of ribosomal and non-ribosomal proteins we showed in Figure 16 at different growth rates. As seen there, the relative value of *J_nr_*(*μ, r*) decreases as *μ* increases, as predicted by the high RBS strength of non-ribosomal proteins. However, the relative value of *Jr*(*μ, r*) increases as *μ* increases. This is possible because the value of *Jr*(*μ, r*) remains mostly constant with respect to *J_nr_*(*μ, r*) (or at least it decreases less than *J_nr_*(*μ, r*) does). Therefore, the relative value of *Jr*(*μ, r*) increases.

In summary, the RBS strength sets the sensitivity of the resources recruitment strength with respect to the flux of free resources. The resources recruitment strength *J_k_*(*μ, r*) of a given protein, related to its expression, can be made dependent on the flux of free resources *μr* (strong RBS) or can be robust with respect to variations of *μr* (weak RBS). This allows to set how much of a given protein (e.g ribosomal or non-ribosomal) will be expressed at different levels of *μr* (i.e. at different growth rates).

This differential expression may have evolved in *E. coli* and other micro-organisms to encode the mass distribution of ribosomal and non-ribosomal proteins as shown in Figure 16 (right). The cell achieves a fairly constant absolute expression of non-ribosomal proteins by using a high RBS strength to express them. This way, as the growth rate increases, (tantamount to *μr*) the value of *J_nr_*(*μ, r*) will decrease, compensating the increase of expression induced by the increase of the total cell protein mass. On the other hand, the cell uses much weaker RBSs to express the ribosomal proteins. This way the value of *J_r_*(*μ, r*) will remain mostly constant with respect to *J_nr_*(*μ, r*). As a consequence, the absolute expression of ribosomal proteins increases with growth rate as the value of *J_nr_*(*μ, r*) decreases, causing the fraction of ribosomes to increase.

## 18. Host-circuit interaction shapes the optimal productivity of proteins

There are multiple ways to increase the expression of a given exogenous protein, including the choice of the expression vector, optimizing the use of codons, co-expression of chaperones to aid protein folding, etc. [32]. Here we pay attention to the differential role of RBS and promoter strengths in determining protein abundance. We evaluated the mass and specific productivities across the expression space *N_A_ω_A_*, 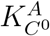 for a given exogenous protein *A* at different values of intracellular substrate availability using the average host dynamics at steady state and the evaluation of protein mass productivity derived in sections 10 and 11.

First we considered the average values of the RBS-strength related parameters 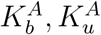 obtained in section 16 for a non-ribosomal protein (see Table 3) and we varied the promoter strength times the gene copy number *N_A_ω_A_* in the range [0.1, 12500] times the average value of the non-ribosomal promoter strengths obtained from the data in [19]. This gives maximum value *N_A_ω_A_* = 375 (mRNA · min^−1^) well within the average transcription rate in *E. coli* if we consider a gene copy number around 100 (e.g. a medium-high copy number plasmid). Figure 19 shows the results obtained. As expected, the cell growth rate abruptly decreases as the mass fraction of the exogenous protein increases for increasing values of *N_A_ω_A_*. Notice that the mass productivity of the exogenous protein reaches a maximum at *N_A_ω_A_* ≈ 30. Thus, for a constitutive promoter with high transcription rate *ω_A_* ≈ 3 (mRNA · min^−1^) the maximum productivity is achieved for a gene copy number *N_A_* ≈ 10 typical of low copy number plasmids. Interestingly, this maximum is independent of the substrate availability, and only depends on the promoter strength and gene copy number.

**Figure 19:**
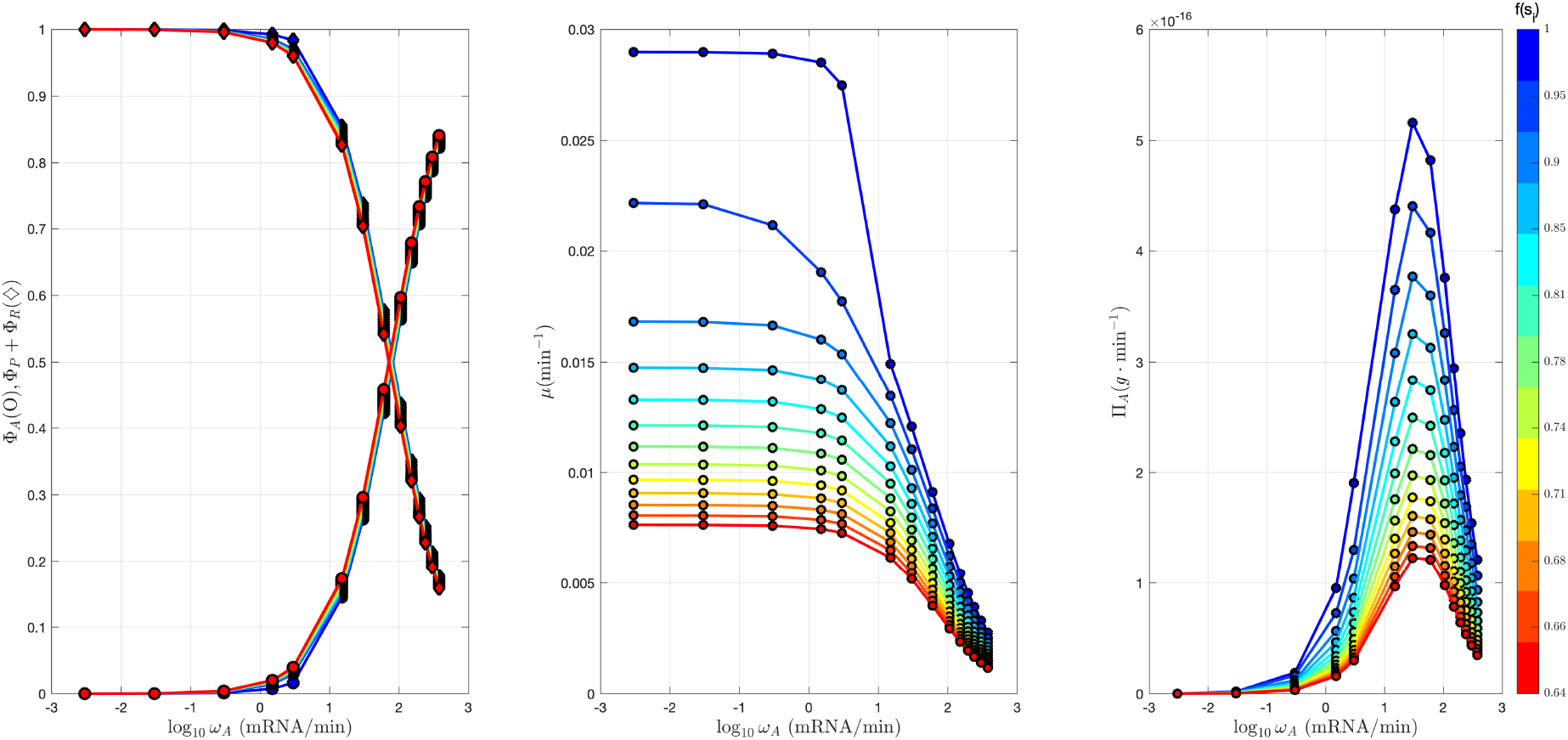
Effect of promoter and gene copy number variation on protein ○ and cell ◊ mass fractions (Left) cell growth rate (Center) and mass productivity (Right) for varying intracellular substrate concentration.

Next we considered two values *N_A_ω_A_* = {3, 150} corresponding to low and high promoter strength times the gene copy number and we varied 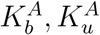 in the ranges considered in section 15 so as to achieve a continuous varying RBS strength 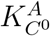. Figure 20 shows the results obtained. Again, the cell growth rate abruptly decreases as the mass fraction of the exogenous protein increases for increasing values of the RBS-strength. Notice though that the main affecting factor is the promoter strength times gene copy number *N_A_ω_A_*. Thus, for low values of *N_A_ω_A_*, increasing RBS strength gives rise to increasing protein mass productivity. Only for high values of *N_A_ω_A_* there appears a maximum of the mass productivity as a function of the RBS strength. Notice also that in this case, the maximum also varies as a function of the substrate availability.

**Figure 20:**
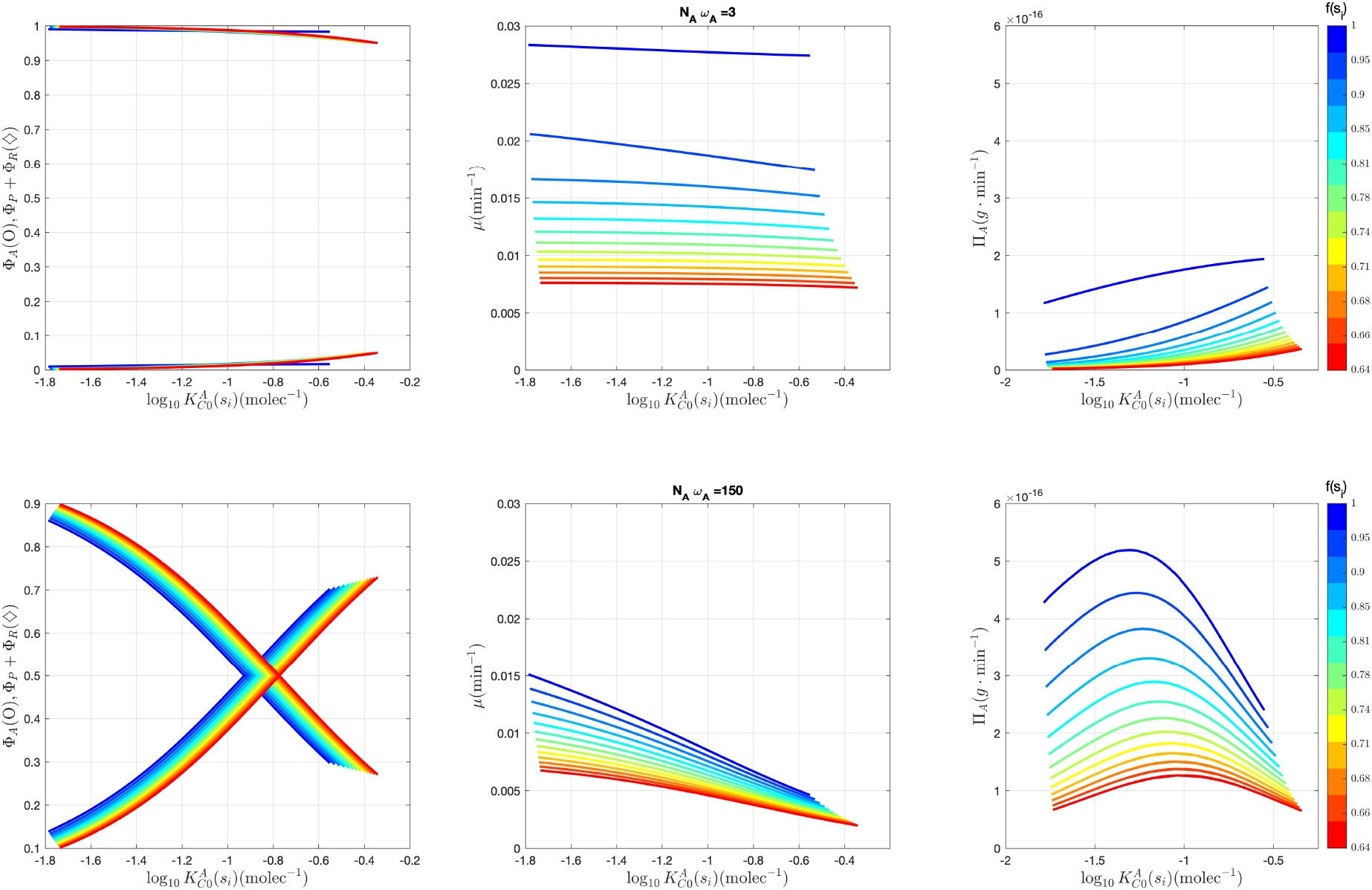
Effect of RBS strength variation on protein and cell mass fractions (Left) cell growth rate (Center) and mass productivity (Right) for varying intracellular substrate concentration and the two values *N_A_ω_A_* = {3, 150}.

Finally, Figure 21 shows the variation of protein mass productivity across the expression space *N_A_ω_A_*, 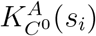 for different values of substrate availability. Notice the optimal productivity subspace corresponds to a hyperbola in the (*N_A_ω_A_*, 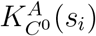)-space showing a trade-off between promoter and RBS strengths. The figure also shows the average values of *N_x_ω_x_*, 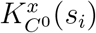 of the *E. coli* host non-ribosomal and ribosomal proteins respectively. Interestingly, ribosomal and non-ribosomal proteins lay in opposite branches of the optimal subspace. While the host ribosomal proteins have evolved a expression strategy based on having a low RBS strength and high promoter strength, the non-ribosomal ones have evolved the opposite strategy. Notice also that while for ribosomal proteins the values of *N_x_ω_x_*, 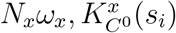 do not appreciably change as a function of substrate, for non-ribosomal proteins the effective RBS strength 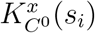 increases as the substrate availability decreases so that their expression keep within the optimal subspace. Therefore, in agreement with our findings in Section 17 the weak promoter-strong RBS strategy versus the strong promoter-weak RBS one is not determined by optimal productivity, but by the strategic decision on the absolute expression should increase with growth rate or not.

**Figure 21:**
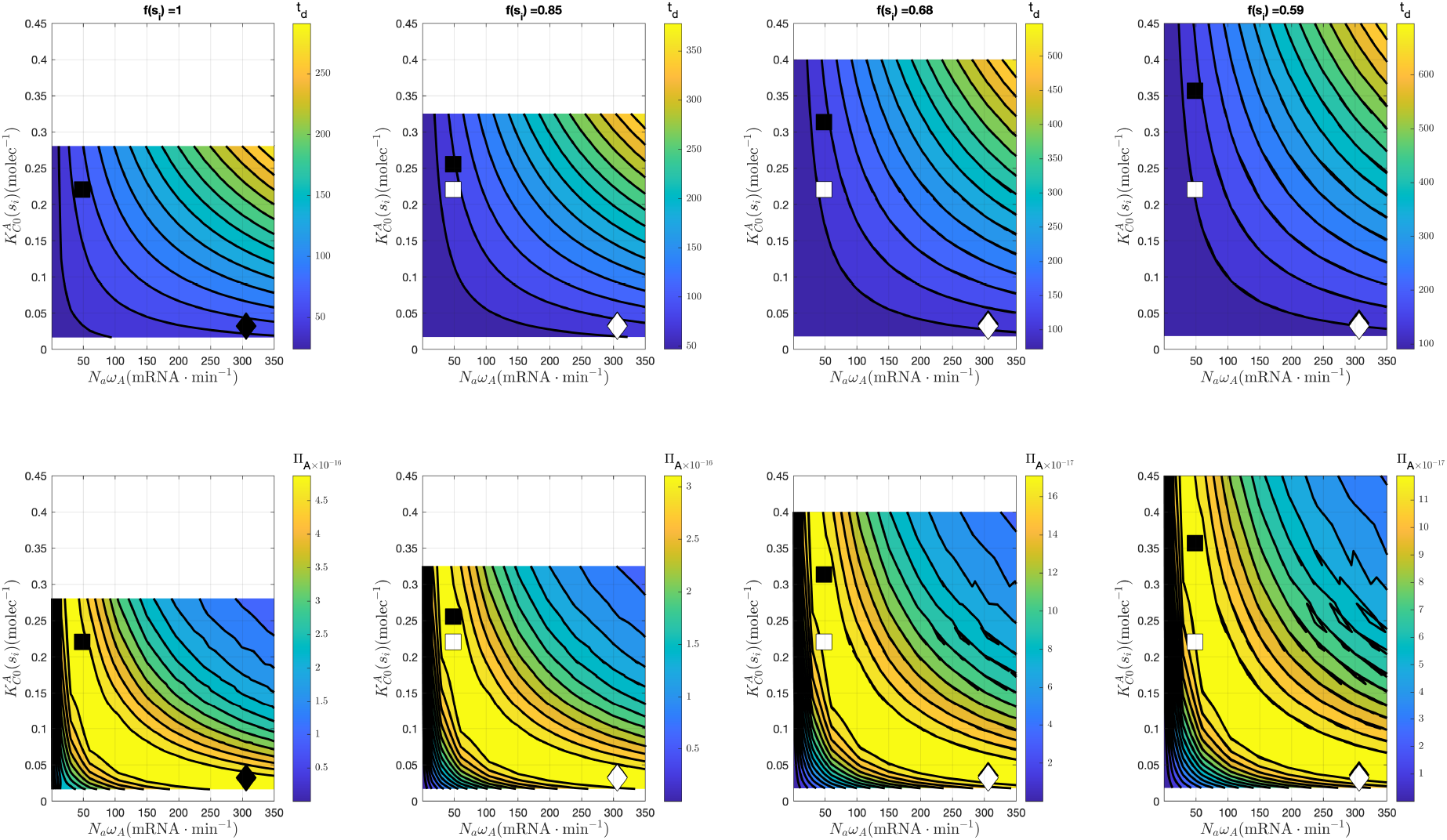
Cell duplication time (top row) and protein mass productivity (bottom row) across the expression space *N_A_ω_A_*, 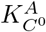 for different values of substrate availability. The black symbols (□, ◊) correspond to the average values of *N_x_ω_x_*, 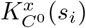 of the *E. coli* host non-ribosomal and ribosomal proteins respectively (see Table 3). The white symbols correspond to the value of *N_x_ω_x_*, 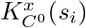 for saturated substrate.

## 19. Conclusions

In this work we have presented a small-size model of gene expression dynamics accounting for host-circuit interactions. The good agreement between the predictions of our model and experimental data highlight the relevance of the cellular resources recruitment strength defined in our model as a key functional coefficient that allows to explain the distribution of resources among the host and the genes of interest and the relationship between the usage of resources, cell growth and protein productivity. This functional coefficient explicitly takes into account the interplay between the flux of available free resources and lab-accessible gene characteristics. In particular, the promoter and RBS strengths.

Though we fitted the model to *E. coli*, our findings can be extrapolated to other microorganisms, and the model can be easily fitted using a small amount of experimental data of the host cell.

Among other predictions, the model provides insights into how the differential role of promoter and RBS strengths in protein expression may have evolved in *E. coli* and other micro-organisms to encode the mass distribution between ribosomal and non-ribosomal proteins as a function of cell growth rate. Weak transcription and strong translation and the complementary strong transcription and weak translation arise a two equally optimal protein productivity strategies in the expression space but with different characteristics from the point of view of the resulting copy number of the expressed protein as a function of growth rate. The capacity of the defined resources recruitment strength functional coefficients to capture the interaction between growth, cell resources and gene expression characteristics is reflected in the fact that we did not estimate their parameters by trying and directly fitting the available experimental relative distribution of resources recruitment strengths. Instead, we sought the model to fit the cell specific growth rate and saw that this implied ribosomal and non-ribosomal resources recruitment strengths such that their relative values fitted the experimental ones.

The model also explains some of the phenomena typically encountered when building protein expression systems in synthetic biology. Thus, for instance, it explains the limited effect that increasing the RBS strength has to increase the expression of a given protein of interest, saturating at high RBS strengths. In this context, our model may also be useful for design purposes in synthetic biology where it can be used to design the proper strategy in the expression space depending on the desired expression behaviour as a function of growth rate.

Further extensions of the model can be easily implemented. Thus, the model explicitly considers the relationship between the cell specific growth rate and the population dynamics, so it can be integrated within a multi-scale framework that considers the macroscopic extracellular dynamics of the substrate and population of cells in a bioreactor. The possibility to consider expression systems using orthogonal ribosomes can also be implemented without much difficulties.

## 20. Acknowledgements

This work was partially supported by grant MINECO/AEI, EU DPI2017-82896-C2-1-R. F.N. Santos-Navarro is grateful to grant PAID-01-2017 (Universitat Politècnica de València)

## 21. Authors contributions

J. Picó conceived the study. J. Picó and F.N. Santos-Navarro designed the methods and experiments. F.N. Santos-Navarro implemented the methods and performed the experiments. J. Picó and F.N. Santos-Navarro analysed the results and wrote the manuscript. J. Picó edited and approved the final manuscript.

